# Deep Boosted Molecular Dynamics (DBMD): Accelerating molecular simulations with Gaussian boost potentials generated using probabilistic Bayesian deep neural network

**DOI:** 10.1101/2023.03.25.534210

**Authors:** Hung N. Do, Yinglong Miao

## Abstract

We have developed a new Deep Boosted Molecular Dynamics (DBMD) method. Probabilistic Bayesian neural network models were implemented to construct boost potentials that exhibit Gaussian distribution with minimized anharmonicity, thereby allowing for accurate energetic reweighting and enhanced sampling of molecular simulations. DBMD was demonstrated on model systems of alanine dipeptide and the fast-folding protein and RNA structures. For alanine dipeptide, 30ns DMBD simulations captured up to 83-125 times more backbone dihedral transitions than 1µs conventional molecular dynamics (cMD) simulations and were able to accurately reproduce the original free energy profiles. Moreover, DBMD sampled multiple folding and unfolding events within 300ns simulations of the chignolin model protein and identified low-energy conformational states comparable to previous simulation findings. Finally, DBMD captured a general folding pathway of three hairpin RNAs with the GCAA, GAAA, and UUCG tetraloops. Based on Deep Learning neural network, DBMD provides a powerful and generally applicable approach to boosting biomolecular simulations. DBMD is available with open source in OpenMM at https://github.com/MiaoLab20/DBMD/.

## Introduction

Molecular dynamics (MD) is a powerful computational technique for simulating biomolecular dynamics at an atomistic level^1^. With recent advances in computing hardware and software developments, timescales accessible to MD simulations have significantly increased^2, 3^. However, conventional MD (cMD) is often limited to tens to hundreds of microseconds^4, 5^ for simulations of typical biomolecular systems, and cannot attain the timescales required to observe many biological processes of interest, which typically occur over milliseconds or longer with high energy barriers (e.g., 8-12 kcal/mol)^6^.

Many enhanced sampling techniques have been developed during the last several decades to overcome the challenges mentioned above^7–11^. In particular, Gaussian accelerated molecular dynamics (GaMD) is an enhanced sampling that technique works by applying a harmonic boost potential to smooth biomolecular potential energy surface^27^. Since this boost potential exhibits a near Gaussian distribution, cumulant expansion to the second order (“Gaussian approximation”) can be applied to achieve proper energetic reweighting^28^. GaMD allows for simultaneous unconstrained enhanced sampling and free energy calculations of large biomolecules ^27^. GaMD has been successfully demonstrated on enhanced sampling of ligand binding, protein folding, protein conformational change, as well as protein-membrane, protein-protein, and protein-nucleic acid interactions^3^. GaMD has been implemented in widely used simulation packages including AMBER^27^, NAMD^29^, OpenMM^30^, GENESIS^31^, and TINKER-HP^32^.

Recently, Machine Learning/Deep Learning techniques (ML/DL) have been combined with MD methods to enhance the sampling of biomolecular simulations. DeepDriveMD is a DL driven adaptive MD method designed specifically to simulate protein folding^33^. In DeepDriveMD, DL was utilized to reduce the dimensionality of MD simulations to automatically build latent representations that correspond to biophysically relevant collective variables (CVs) and drive MD simulations to automatically sample potentially novel conformational states based on the CVs^33^. DeepDriveMD has been demonstrated to speed up the folding simulations of Fs-peptide and the fast-folding variant of the villin head piece protein by at least 2.3 folds^33^. The State Predictive Information Bottleneck (SPIB) approach was applied as a deep neural network to learn a priori CV for well-tempered metadynamics from undersampled trajectories^34^. The well-tempered metadynamics performed along the biased SPIB-learned CVs were shown to achieve > 40 times acceleration in simulating the left- to right-handed chirality transitions in a synthetic helical peptide and permeation of a small benzoic acid molecule through a synthetic, symmetric phospholipid bilayer^34^. Moreover, denoising diffusion probabilistic models were combined with replica exchange MD to achieve superior sampling of biomolecular energy landscape at temperatures that were not simulated without the assumption of particular slow degrees of freedom^35^. The temperature was treated as a fluctuating random variable and not a control parameter to allow for the direct sampling from the joint probability distribution in configuration and temperature space. The procedure was shown to discover transition and metastable states that were previously unseen at the temperature of interest and bypass the need to perform simulations for a wide range of temperatures^35^.

In this work, we have developed a new Deep Boosted Molecular Dynamics (DBMD) method. In DBMD, probabilistic Bayesian neural network models were used to construct boost potentials that exhibit Gaussian distribution with minimized anharmonicity for accurate energetic reweighting and enhanced sampling. DBMD has been demonstrated on model systems of the alanine dipeptide in explicit and implicit solvent, the chignolin fast-folding protein, and three hairpin RNAs with the GCAA, GAAA, and UUCG tetraloops.

## Methods

### Theory of DBMD

In DBMD, boost potentials Δ*V* are optimized using DL to follow Gaussian distribution with minimized anharmonicity. Considering a system comprised of *N* atoms with coordinates 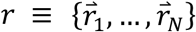 and momenta 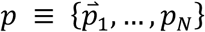 the system Hamiltonian can be expressed as:

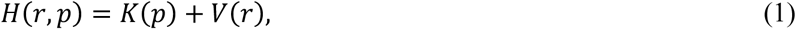

where *K*(*p*) and *V*(*r*) are the system kinetic and total potential energies, respectively. To enhance biomolecular conformational sampling, boost potentials can be added to the system potential energies. According to the DBMD algorithm, the boost potential can be calculated as the following^27^:

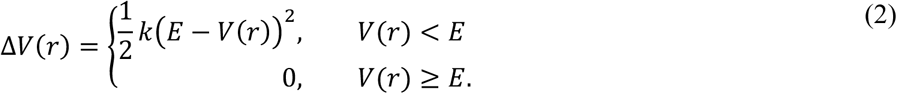

where *E* is the reference energy for adding boost potential and *k* is the harmonic force constant. Here, the reference energy can be set in a range: 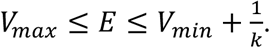 The harmonic force constant is calculated as 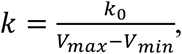 with the effective harmonic force constant (*k*_0_ ∈ (0,1]. Accordingly, the reference energy can be expressed as 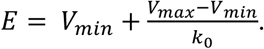 Here, *E* = *V_max_* when *k*_0_ = 1, and the smaller the *k*_0_ values, the higher the reference energy *E*. In DBMD, we introduce a parameter called the reference energy factor (*η*) valued between 0 and 1 to avoid exceedingly large *E* and control the acceleration during simulations. Physically, *V*_max_ + *η* ∗ |*V*_*max*_| represents the upper limit of the reference energy *E*.

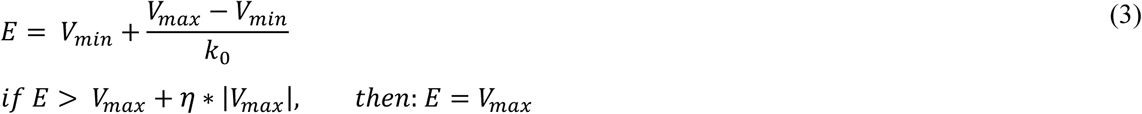

Therefore, the boost potential can be rewritten as:

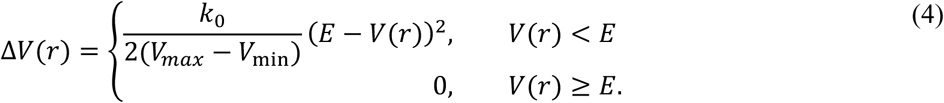

To characterize the extent to which Δ*V* follows a Gaussian distribution, its distribution anharmonicity *γ* is calculated as:

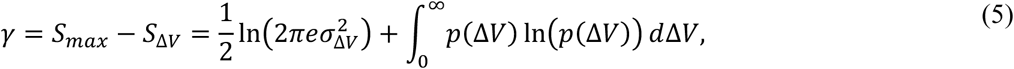

where Δ*V* is dimensionless as divided by *k*_*B*_*T* with *k*_*B*_ and *T* being the Boltzmann constant and system temperature, respectively, and 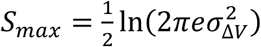 is the maximum entropy of Δ*V*^28^. When *γ* is zero, Δ*V* follows exact Gaussian distribution with sufficient sampling. Reweighting by approximating the ensembled-averaged Boltzmann factor with cumulant expansion to the 2^nd^ order (“Gaussian approximation”) can accurately recover the original free energy landscape^27, 36^. As *γ* increases, the Δ*V* distribution becomes less harmonic, and the reweighted free energy profile obtained from cumulant expansion to the 2^nd^ order would deviate from the original^27^. The anharmonicity of Δ*V* distribution serves as an indicator of the enhanced sampling convergence and accuracy of the reweighted free energy^27^.

### Deep Learning of Potential Energies

In DBMD, the probabilistic Bayesian neural network model within the *TensorFlow Probability*^37^ module was applied to minimize the anharmonicity of boost potentials Δ*V*. The probabilistic model was initiated with the definition of a prior distribution for weights. A standard normal distribution was adopted as the prior distribution since the central limit theorem asserts that a properly normalized sum of samples will approximate a normal distribution^38, 39^. Here, a multivariate normal distribution with a diagonal covariance matrix was used, with the mean values initialized to zero and the variances 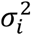 to one^38^.

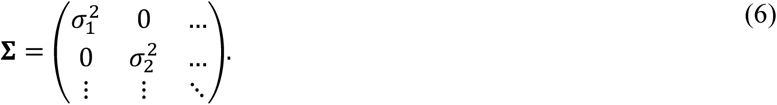

The posterior distribution was also set to be a multivariate Gaussian distribution, but the off-diagonal elements in the covariance matrix were allowed to be non-zero. This was achieved with a lower-triangular matrix **L** with positive-valued diagonal entries such that **Σ** = **LL**^T^, and the triangular matrix can be obtained through Cholesky decomposition of the covariance matrix^38^.

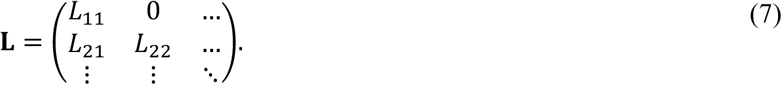

Finally, the probabilistic layers were defined using the *DenseVariational* function of the *TensorFlow Probability* module^37–39^. Our Bayesian neural network model consisted of two or four dense variational layers of two different types, namely *L1* and *L2*. The first dense variational layer *L1* had 64 filters, with a sigmoid activation function to enable the fitting of non-linear data^38, 39^. The second dense variational layer *L2* used the *IndependentNormal*^37^ function to parameterize a normal distribution and capture aleatoric uncertainty, with an event shape equal to one^38, 39^. The prior and posterior distributions used in both *L1* and *L2* were specified above. Testing simulations have showed us that the number of the second dense variational layer *L2* could significantly affect the average and standard deviation of the output boost potentials after DL. Overall, the lower the numbers of *L2*, the wider the distributions and the higher the average boost potentials. Therefore, to balance between the stability and sampling of the simulations as well as the learning speed, we included one *L2* layer in the DL model for explicit-solvent simulations and three *L2* layers for implicit-solvent simulations. The input and output shape were set to one since both the potential energies and boost potentials were scalars.

### Workflow of DBMD

The workflow of DBMD is shown in Figure 1. First, a short cMD was performed on the biological system of interest, and the potential statistics (*V_min_* and *V_max_*) were collected as parameters for pre-equilibration of DBMD simulation. During the pre-equilibration, the effective harmonic force constants (*k_0P_* and *k_0D_*) were kept fixed at (1.0, 1.0) for explicit-solvent simulations and (0.05, 1.0) for implicit-solvent simulations. The boost potentials were calculated based on equation (4), and the potential statistics (*V_min_* and *V_max_*) were updated during pre-equilibration. The system total and dihedral potential energies from the pre-equilibration were then collected (Figure 1a), which served as the *X* inputs for the probabilistic Bayesian DL models^37, 38^ (Figure 1b). Initial boost potentials were randomly generated from the system potential energies and randomly assigned *k_0_* using equations (3-4) and used as the *Y* inputs for DL (Figure 1c). DL was carried out in multiple iterations until the output boost potentials followed Gaussian distribution with anharmonicity *γ* < 0.01 (Figure 1d). If *γ* ≥ 0.01, the generated boost potentials were used as *Y* inputs to retrain the DL model until *γ* < 0.01 (Figure 1e). Based on the potential statistics learnt until the last frame of the pre-equilibration (*V_min_*, *V_max_*, *V*, and Δ*V*), the effective harmonic force constants were calculated as following:

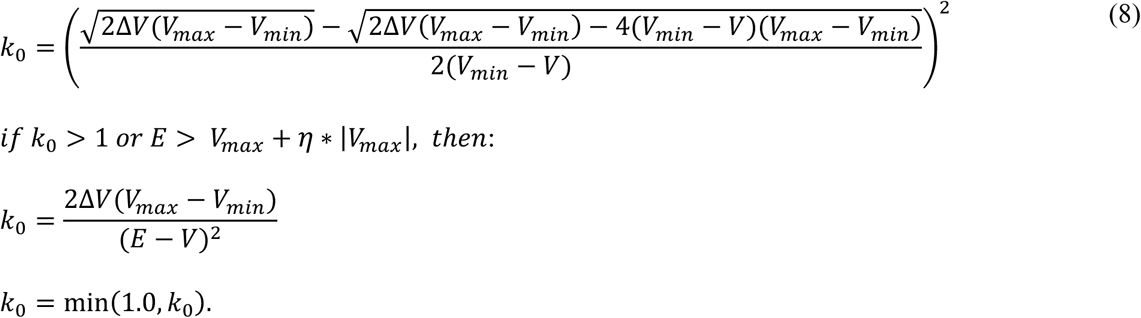

**Figure 1.**
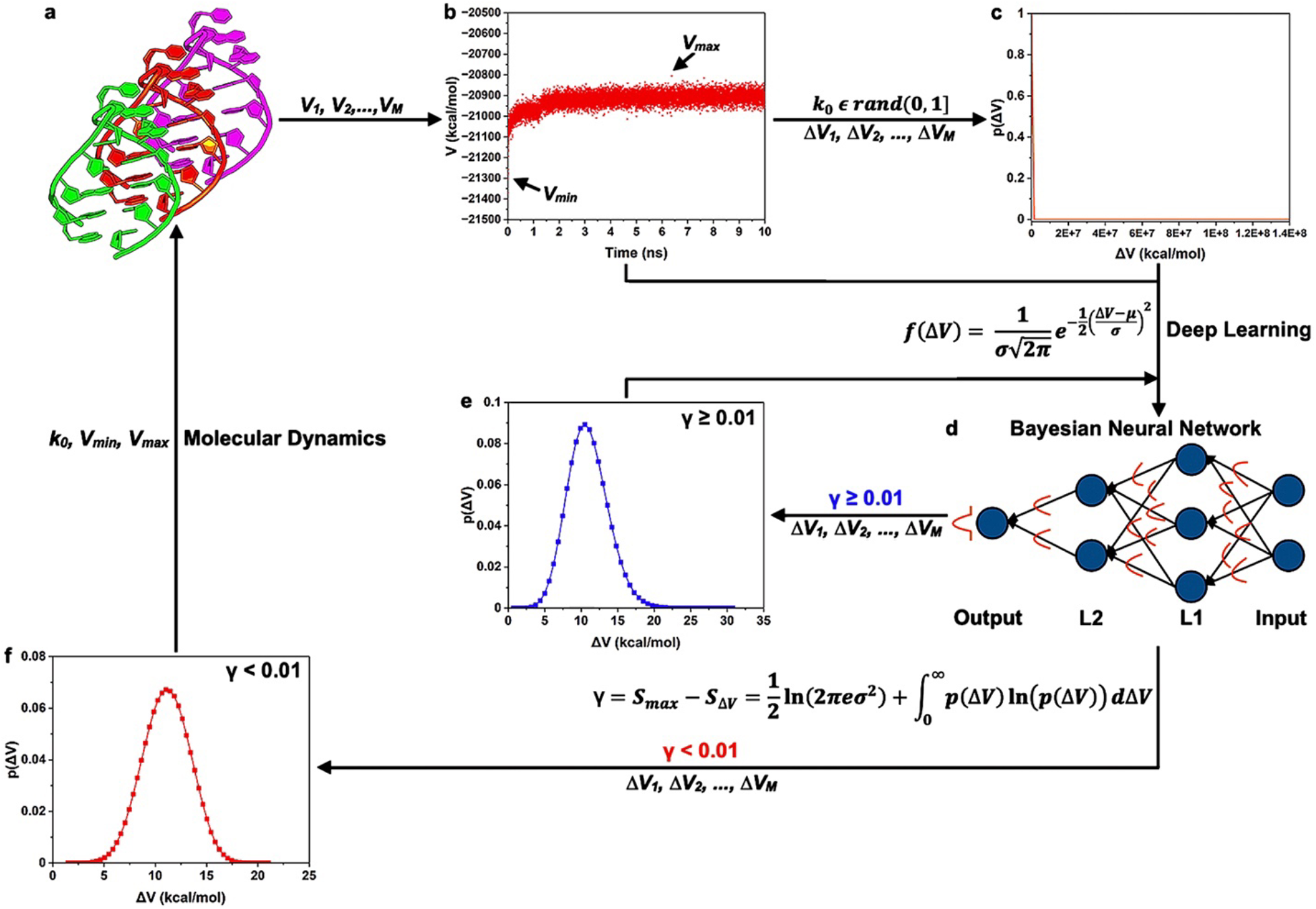
Summary of Deep Boosted Molecular Dynamics (DBMD) **(a)** First, molecular dynamics (MD) simulation is performed on the system of interest. **(b)** The system potential energies from finished simulation frames (*V_1_, V_2_, …, V_M_*) are collected as the *X* inputs for the probabilistic Bayesian Deep Learning (DL) model. **(c)** Reference boost potentials (Δ*V_1_, ΔV_2_, …, ΔV_M_*) were generated from the collected system potential energies and randomized effective harmonic force constants *k_0_* to serve as the *Y* inputs for the DL. **(d)** The probabilistic Bayesian neural network was trained to generate boost potentials that follow Gaussian distribution with the probability density function *f*(Δ*V*). Here, Δ*V* is boost potential, and *μ* and *σ* are the average and standard deviation of the boost potentials. DL is carried out in multiple iterations until the anharmonicity of output boost potentials *γ* < 0.01. **(e)** If the anharmonicity of output boost potential *γ* is ≥ 0.01, the generated boost potentials are used as *Y* inputs to retrain the DL model until *γ* < 0.01. **(f)** Finally, the effective harmonic force constants *k_0_* are calculated from the system potential energy (*V_M_*) and used as input alongside the minimum and maximum of potential energy (*V_min_* and *V_max_*) **(b)** for the next round of enhanced sampling simulation.

and used as input alongside *V_min_* and *V_max_* to equilibrate the simulation system (Figure 1f). The equilibration usually consisted of multiple rounds, with the effective harmonic force constants (*k_0P_* and *k_0D_*) kept fixed and potential statistics (*V_min_* and *V_max_*) updated in each round. DL was carried out at the end of each round using the updated potential energies as inputs, with the same DL model as obtained at the end of the pre-equilibration (Figure 1). Finally, the effective harmonic force constants (*k_0P_* and *k_0D_*) and potential statistics (*V_min_* and *V_max_*) taken from the last round of the equilibration were used as input parameters for DBMD production simulations (Figure 1f), during which the effective harmonic force constants and potential statistics were kept fixed, and boost potentials were calculated based on equation (4).

### System Setup and Simulation Protocols

Simulations of the alanine dipeptide and chignolin were performed using the AMBER ff99SB force field parameter set^40–43^. The LEaP module in the AmberTools package^40–43^ were used to build the simulation systems. For the DBMD simulations in explicit solvent, alanine dipeptide was solvated in a TIP3P^44^ water box that extended ∼8 Å from the solute surface. The unfolded chignolin with a sequence of 10 residues (GYDPETGTWG)^45^ was solvated in a TIP3P^44^ water box that extended ∼10 Å from the solute surface. The final system for alanine dipeptide in explicit solvent, alanine dipeptide in implicit solvent, and chignolin in explicit solvent contained 1912, 22, and 6773 atoms, respectively.

Simulations of the hairpin RNAs with the GCAA, GAAA, and UUCG tetraloops were carried out using the AMBER Shaw force field parameter set^46^, starting from their unfolded states. The sequences of the hairpin RNAs with GCAA, GAAA, and UUCG tetraloops were GGGCGCAAGCCU (12 nucleotides)^47^, CGGGGAAACUUG (12 nucleotides)^48^, and GGCACUUCGGUGCC (14 nucleotides)^49^, respectively. The final systems of the hairpin RNAs with GCAA, GAAA, and UUCG tetraloops in implicit solvent contained 389, 390, and 447 atoms, respectively. All simulations were carried out at 300K temperature.

For the explicit-solvent simulations, periodic boundary conditions were applied, and bonds containing hydrogen atoms were restrained with the SHAKE^50^ algorithm. Weak coupling to an external temperature and pressure bath was necessary to control both temperature and pressure^51^. The electrostatic interactions were calculated using the particle mesh Ewald (PME) summation^52^ with a cutoff of 8.0-9.0 Å for long-range interactions. For the implicit-solvent simulations, the generalized Born solvent model 2 (GBn2)^53^ parameters were used. No nonbonded cutoff was set and no periodic boundary condition was used in the implicit-solvent simulations. The solute and solvent dielectric constants were set to 1.0 and 78.5, respectively, and the effect of a non-zero salt concentration was achieved by setting the Debye-Huckel screening parameter^54^ to 1.0/nm. A 2-fs timestep with the SHAKE^50^ algorithm applied was used in all simulations.

For alanine dipeptide, the simulations consisted of a 2ns short cMD, followed by a 2ns DBMD pre-equilibration, one round of 2ns DBMD equilibration, and three independent 30ns DBMD production simulations. The reference energy factors were set to zero for both total and dihedral potential energy (*η*_0_ and *η*_1_), i.e., *E = V_max_*. For chignolin, the simulation involved a 5ns cMD, a 2ns DBMD pre-equilibration, two rounds of 5ns DBMD equilibration, and three independent 300ns DBMD production simulations, with *η*_0_ and *η*_1_ both set to 0.05. For the hairpin RNAs with GCAA, GAAA, and UUCG tetraloops, the simulations consisted of a 20ns cMD, followed by a 5ns DBMD pre-equilibration, three rounds of 5ns DBMD equilibration, and three-four independent 2µs DBMD production simulations. *η*_0_ and *η*_1_ were set to 0.05 and 0.05 for GCAA, 0.05 and 0.0 for GAAA, and 0.0 and 0.0 for UUCG RNA tetraloops. The simulation frames were saved every 0.1 ps. The CPPTRAJ^55^ tool was used for simulation trajectory analysis.

Finally, the PyReweighting toolkit^28^ was used to compute the potential of mean force (PMF) profiles of the backbone dihedrals Phi and Psi (Φ and Ψ) in the alanine dipeptide (Figure 2a). The C_α_-atom root-mean-square deviation (RMSD) of residues Y2-W9 of chignolin relative to the 1UAO^45^ PDB and C_α_-atom radius of gyration (Rg) of residues Y2-W9 were selected as RCs to calculate the PMF profiles in the simulations of chignolin folding. The heavy-atom RMSD of the whole hairpin RNAs with tetraloops relative to respective PDB structures (1ZIH^47^ for GCAA, 2ADT^48^ for GAAA, and 2KOC^49^ for UUCG) and the G1-U12, C1-G12, and G1-C14 center-of-mass (COM) distances were used as RCs to calculate the PMF profiles in the simulations of hairpin RNAs with tetraloops. A bin size of 6°, 1.0 Å, and 1.0-2.0 Å and cutoff of 10, 100, and 100-500 in one bin were used for reweighting of DBMD simulations of alanine dipeptide, chignolin, and hairpin RNAs with tetraloops, respectively.

**Figure 2.**
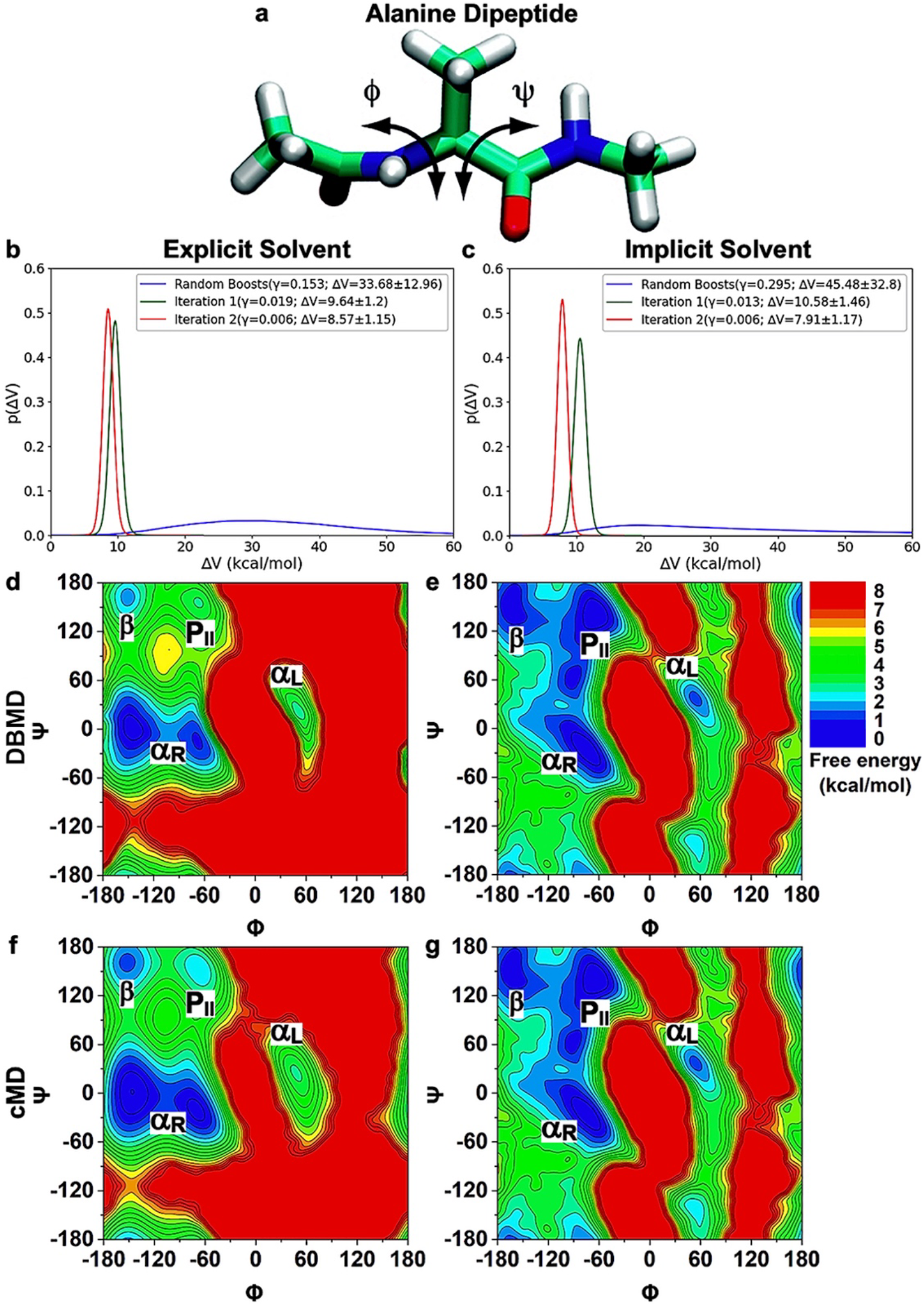
DBMD simulations of alanine dipeptide. **(a)** Schematic representation of backbone dihedrals Phi (Φ) and Psi (Ψ) dihedrals of alanine dipeptide. **(b-c)** Representative distributions of randomly generated dual boost potentials and DL-generated boost potentials iterated until *γ* < 0.01 from the potential energies collected from the pre-equilibration of the alanine dipeptide in explicit solvent **(b)** and implicit solvent **(c)**. The legends include the anharmonicity and average ± standard deviation of the dual boost potentials. **(d-g)** 2D Potential of mean force (PMF) free energy profile of backbone dihedrals (Φ, Ψ) of alanine dipeptide calculated from three 30ns DBMD simulations **(d-e)** compared to 1μs cMD simulations **(f-g)** in explicit solvent **(d, f)** and implicit solvent **(e, g)**. The low-energy states are labeled corresponding to the right-handed α helix (α_R_), left-handed α helix (α_L_), π-sheet (π), and polyproline II (P_II_) conformations.

## Results

### DBMD Simulations of Alanine Dipeptide

DBMD simulations were performed on alanine dipeptide on alanine dipeptide (Figure 2a) in explicit and implicit solvent. Representative distributions of randomly generated boost potentials and the boost potentials generated by DL for alanine dipeptide in explicit and implicit solvent are shown in Figure 2b and 2c, respectively. DL was able to reduce the anharmonicity from 0.153 for the randomly generated boost potentials to 0.019 and 0.006 in two iterations of the explicit-solvent simulation (Figure 2b), and from 0.295 to 0.013 and 0.006 in two iterations of the implicit-solvent simulation (Figure 2c).

The time courses of the effective harmonic force constants (*k_0P_* and *k_0D_*) as well as the total and dihedral boost potential parameters (*V_min_*, *V_max_*, and *E*) during the equilibration of the alanine dipeptide in explicit and implicit solvent are shown in **Figure S1**. During the one round of 1ns DBMD equilibration in explicit solvent, the total and dihedral effective harmonic force constants *k_0P_* and *k_0D_* stayed at 0.35 and 1.0, respectively (**Figure S1a**). The minimum total and dihedral potential energies *V_minP_* and *V_minD_* also remained constant at −5,966.96 kcal/mol and 5.92 kcal/mol, respectively (**Figure S1b-S1c**). However, the maximum total and dihedral potential energy *V_maxP_* and *V_maxD_* increased from −5,742.44 kcal/mol and 25.18 kcal/mol to −5,690.89 kcal/mol and 33.16 kcal/mol, respectively (**Figure S1b-S1c**). The reference total and dihedral potential energy for applying boosts were the same as the maximum potential energies. The effective harmonic force constants as well as extrema and reference potential energies in the implicit-solvent equilibration followed similar trends as the explicit-solvent simulation (**Figure S1e-S1g**).

Three independent 30ns DBMD simulations of alanine dipeptide in both explicit and implicit solvent captured more dihedral transitions compared to 1µs cMD simulations (**Figure S2**). In particular, DBMD sampled ∼15, ∼14, and ∼10 Φ dihedral transitions during the 30ns of Sim1, Sim2, and Sim3, respectively, compared to only ∼4 dihedral transitions observed in the 1µs cMD of alanine dipeptide in explicit solvent (**Figure S2a-S2d**). In the implicit-solvent simulations, Sim1, Sim2, and Sim3 sampled ∼17, ∼28, and ∼28 Φ dihedral transitions during the 30ns simulations, respectively, compared to the ∼26 Φ dihedral transitions observed in the 1µs cMD simulation (**Figure S2e-S2h**). Therefore, DBMD accelerated the explicit-solvent simulations by ∼83-125 times and implicit-solvent simulations by ∼22-36 times. Furthermore, the boost potentials applied in DBMD simulations of alanine dipeptide followed Gaussian distributions, with low anharmonicity of 6.2 × 10^−3^ in the explicit-solvent and 1.7 × 10^−4^ in implicit-solvent simulations (**Figure S3a-S3b**). The averages and standard deviations of the added boost potentials were recorded to be 11.2 ± 2.8 and 11.3 ± 2.3 kcal/mol in the explicit and implicit solvent simulations, respectively.

The PMF free energy profiles of alanine dipeptide were calculated for the Φ and Ψ dihedral angles. The 1D PMF free energy profiles were in excellent agreement between DBMD and cMD for both Φ and Ψ in explicit and implicit solvent (**Figure S3c-S3f**). Moreover, the 2D PMF free energy profiles of the (Φ, Ψ) backbone dihedrals showed high degrees of similarity between DBMD and cMD simulations (Figure 2d-2g). In particular, DBMD simulations in explicit solvent sampled five different low-energy conformational states of alanine dipeptide, which centered around (−150°, 159°) in the ϕ3-sheet, (−72°, 162°) in the polyproline II (P_II_), (48°, 18°) in the left-handed α helix (α_L_), and (−148°, 0°) and (−69°, −17°) in the right-handed α helix (α_R_) conformation (Figure 2d). In implicit solvent, DBMD also identified five low-energy conformational states of alanine dipeptide, including ϕ3-sheet centered around (−160°, 150°), P_II_ around (−62°, 140°) and (−90°, 61°), α_L_ around (56°, 34°), and α_R_ around (−70°, −27°) (Figure 2e). The 1D and 2D free energy profiles of (Φ, Ψ) calculated from DBMD simulations were in excellent agreements with previous GaMD simulations performed by AMBER^27^, NAMD^29^, and OpenMM^30^. Therefore, simulations of alanine dipeptide have demonstrated the enhanced sampling capability as well as accuracy of DBMD for both explicit and implicit solvent systems.

### DBMD Simulations of Chignolin Folding

Representative distributions of randomly generated dual boost potentials and the boost potentials generated by DL for chignolin folding are shown in Figure 3a. With the use of DL, the anharmonicity reduced from 0.17 for the randomly generated boost potentials to 0.01 and 0.005 in two iterations (Figure 3a).

**Figure 3.**
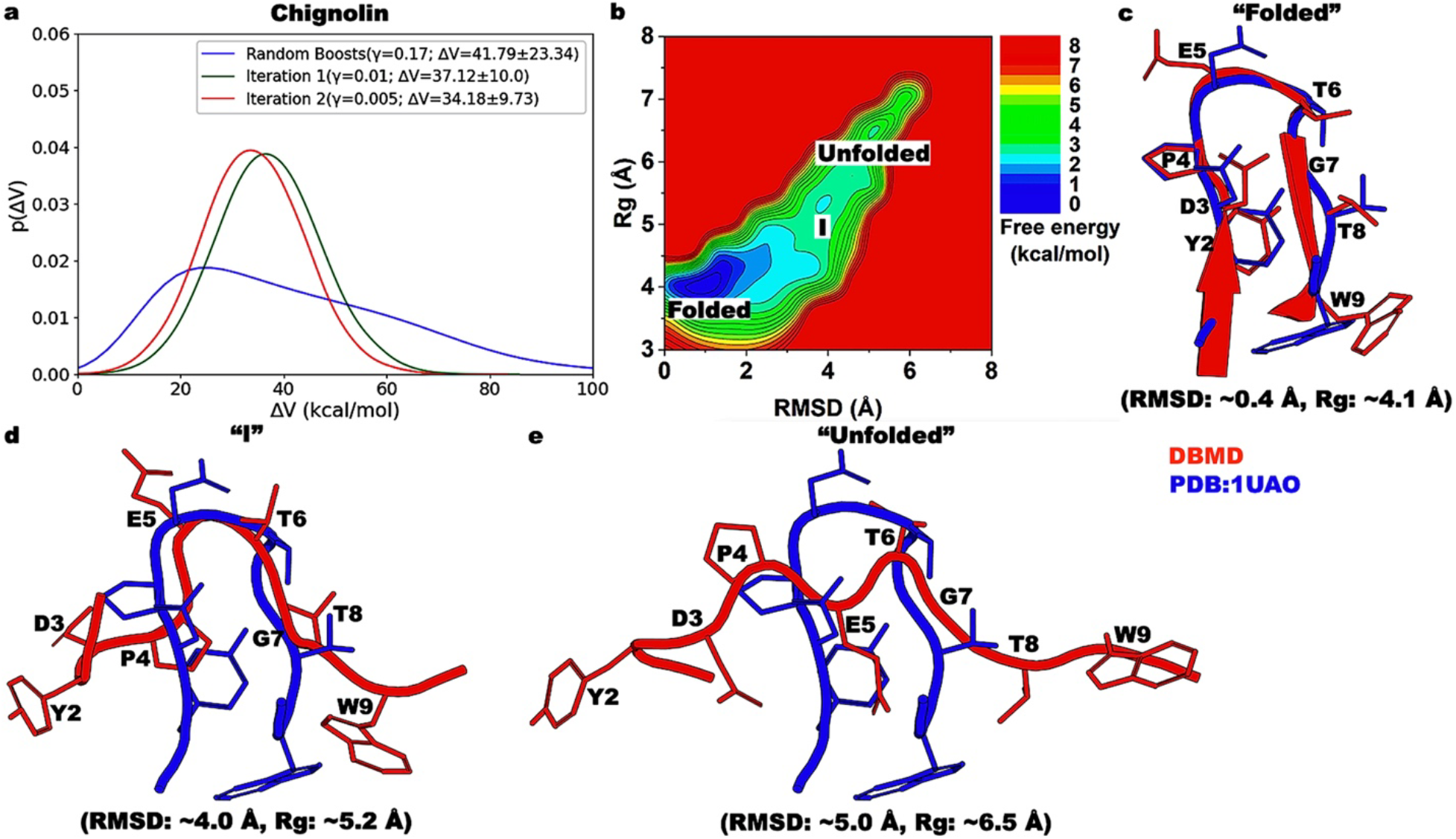
Folding of chignolin in explicit solvent captured by DBMD. **(a)** Representative distributions of randomly generated dual boost potentials and DL-generated boost potentials iterated until *γ* < 0.01 from the potential energies collected from the pre-equilibration of chignolin. The legends include the anharmonicity and average ± standard deviation of the dual boost potentials. **(b)** 2D PMF free energy profile of the C_α_-atom root-mean-square deviation (RMSD) of residues Y2-W9 of chignolin relative to the 1UAO PDB and C_α_-atom radius of gyration (Rg) of residues Y2-W9. The low-energy conformational states are labeled “Folded”, “I”, and “Unfolded”. **(c)** The “Folded” low-energy conformational state compared to the 1UAO PDB structure, for which the RMSD is ∼0.4 Å and the Rg is ∼4.1 Å. **(d)** The intermediate “I” low-energy conformational state compared to the 1UAO PDB structure, for which the RMSD is ∼4.0 Å and the Rg is ∼5.2 Å. **(e)** The “Unfolded” low-energy conformational state compared to the 1UAO PDB structure, for which the RMSD is ∼5.0 Å and the Rg is ∼6.5 Å. The low-energy conformational states are colored red, and the 1UAO PDB structure is colored blue.

The time courses of the effective harmonic force constants (*k_0P_* and *k_0D_*) as well as the total and dihedral boost potential parameters (*V_min_*, *V_max_*, and *E*) during the equilibration of the chignolin fast-folding protein in explicit solvent are shown in **Figure S4**. During the two rounds of 5ns DBMD equilibration, the dihedral effective harmonic force constant *k_0D_* remained at 1.0, while the total effective harmonic force constant *k_0P_* decreased from 0.94 in round one to 0.89 in round two (**Figure S4a**). The minimum total potential energy *V_minP_* increased from −21,388.36 kcal/mol in round one to −20,761.33 kcal/mol in round two (**Figure S4b**). The maximum total potential energy *V_maxP_* increased from −20,742.03 kcal/mol to −20,234.95 kcal/mol and −19,671.23 kcal/mol at the end of round one and two, respectively (**Figure S4b**). The minimum dihedral potential energy *V_minD_* increased from 87.50 kcal/mol in round one to 94.88 kcal/mol in round two (**Figure S4c**). The maximum dihedral potential energy *V_maxD_* increased from 120.52 kcal/mol to 139.37 kcal/mol at the end of round one and 143.53 kcal/mol at the end of round two (**Figure S4c**). While the reference dihedral potential energy *E_D_* was identical to the maximum dihedral potential energy *V_maxD_*, the reference total potential energy *E_P_* was slightly higher than the maximum total potential energy *V_maxP_* (**Figure S4b**).

Three independent 300ns DBMD simulations of chignolin in explicit solvent starting from its extended conformation were able to capture multiple folding and unfolding events of the protein (**Figure S5**). In particular, six, seven, and ten different folding-unfolding events were sampled in Sim1, Sim2, and Sim3 of chignolin (**Figure S5a**). Here, chignolin was considered folded if the C_α_-atom RMSD of residues Y2-W9 was ≤ 1.0 Å. Furthermore, the boost potentials applied in DBMD simulations of chignolin followed the Gaussian distribution, with an anharmonicity of 7.1 × 10^−3^ (**Figure S5c**) and an average of 23.1 ± 5.1 kcal/mol.

The 2D PMF free energy profile of chignolin folding was calculated using the C_α_-atom RMSD relative to the 1UAO^45^ PDB structure and Rg of residues Y2-W9 as RCs. Three different low-energy conformational states of chignolin were identified from the free energy profile, namely “Folded”, intermediate “I”, and “Unfolded” (Figure 3b). The “Folded” low-energy conformational state of chignolin centered around 0.4 Å and 4.1 Å of RMSD and Rg, respectively. In this state, terminal residues Y2-D3 formed β-sheets with residues G7-W9 of chignolin, while the loop formed by the backbone atoms of residues P4-T6 closely matched with the 1UAO^45^ PDB structure (Figure 3c). In the intermediate “I” low-energy conformational state, the C_α_-atom RMSD and Rg were ∼4.0 Å and ∼5.2 Å. Transitioning from the “Folded” to intermediate “I” state, the β-strands were broken apart due to the opposite movement of residues G1-D3 and T8-G10. However, the core loop of chignolin was somewhat maintained with the hydrophilic side chains of residues E5-T6 exposed to the solvent (Figure 3d). Finally, in the “Unfolded” low-energy conformational state, chignolin was fully extended with all amino acids exposed to the solvent, resulting in a RMSD of ∼5.0 Å and Rg of ∼6.5 Å (Figure 3e).

### DBMD Simulations of RNA Folding with Tetraloops

Representative distributions of randomly generated dual boost potentials and the boost potentials generated by DL for the hairpin RNAs with the GCAA, GAAA, and UUCG tetraloops are shown in Figures 4a-6a, respectively. With the use of DL, the anharmonicity reduced from 0.135 for the randomly generated boost potentials to 0.016, 0.015, and 0.009 in three iterations of the GCAA RNA system simulation (Figure 4a). For GAAA, DL lowered the anharmonicity from 0.137 for the random boost potentials to 0.012, 0.01, and 0.008 in three iterations (Figure 5a). For UUCG, the anharmonicity reduced from 0.147 to 0.014 to 0.013 and 0.008 (Figure 6a).

**Figure 4.**
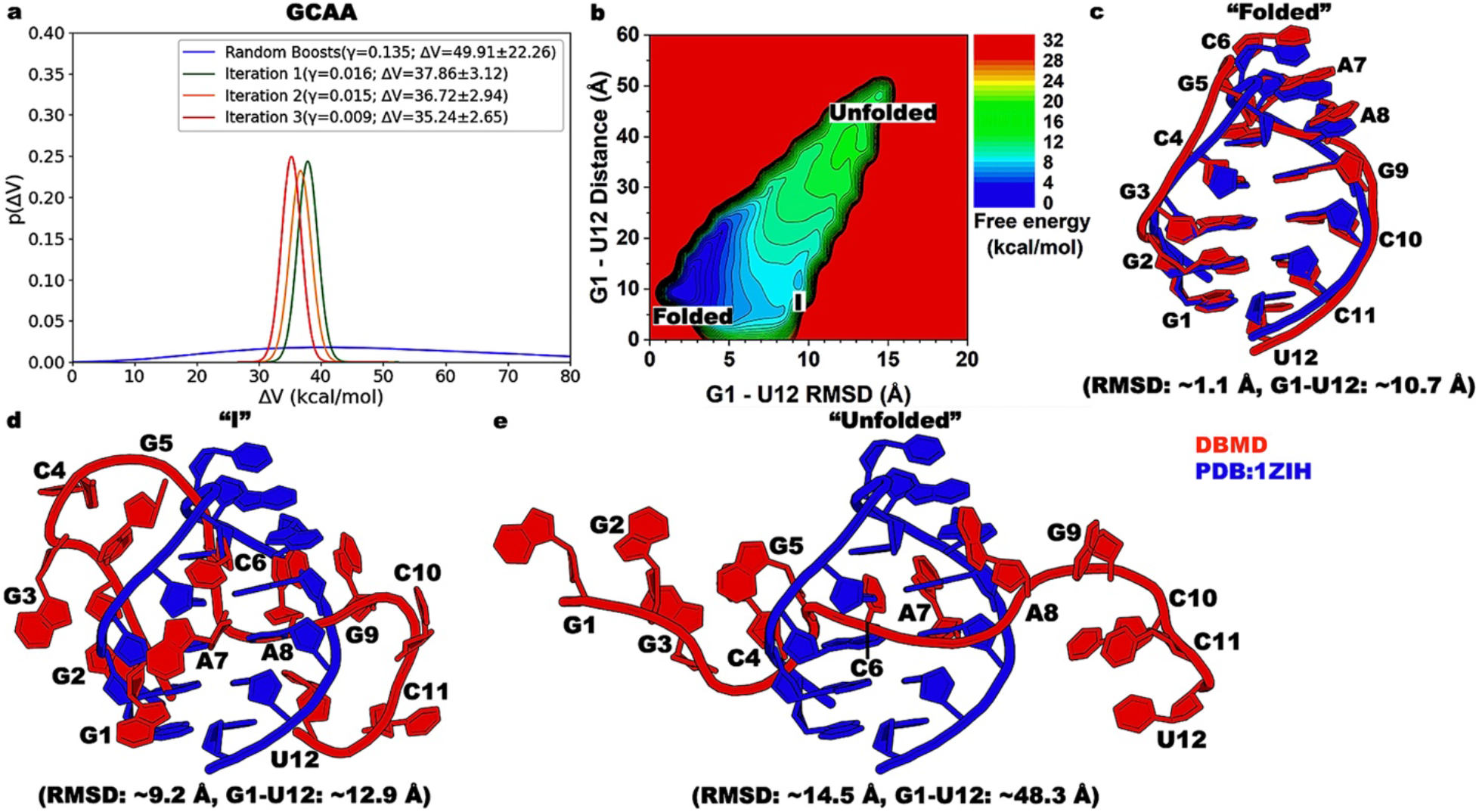
Folding of the 12-mer hairpin RNA with GCAA tetraloop in implicit solvent captured by DBMD. **(a)** Representative distributions of randomly generated dual boost potentials and DL-generated boost potentials iterated until *γ* < 0.01 from the potential energies collected from the pre-equilibration of the 12-mer hairpin RNA with GCAA tetraloop. The legends include the anharmonicity and average ± standard deviation of the dual boost potentials. **(b)** 2D PMF free energy profile of the heavy-atom RMSD of the 12-mer hairpin RNA relative to the 1ZIH PDB and the center of mass (COM) distance between terminal nucleotides G1 and U12. The low-energy conformational states are labeled “Folded”, “I”, and “Unfolded”. **(c)** The “Folded” low-energy conformational state compared to the 1ZIH PDB structure, for which the RMSD is ∼1.1 Å and the G1-U12 distance is ∼10.7 Å. **(d)** The intermediate “I” low-energy conformational state compared to the 1ZIH PDB structure, for which the RMSD is ∼9.2 Å and the G1-U12 distance is ∼12.9 Å. **(e)** The “Unfolded” low-energy conformational state compared to the 1ZIH PDB structure, for which the RMSD is ∼14.5 Å and the G1-U12 distance is ∼48.3 Å. The low-energy conformational states are colored red, and the 1ZIH PDB structure is colored blue.

**Figure 5.**
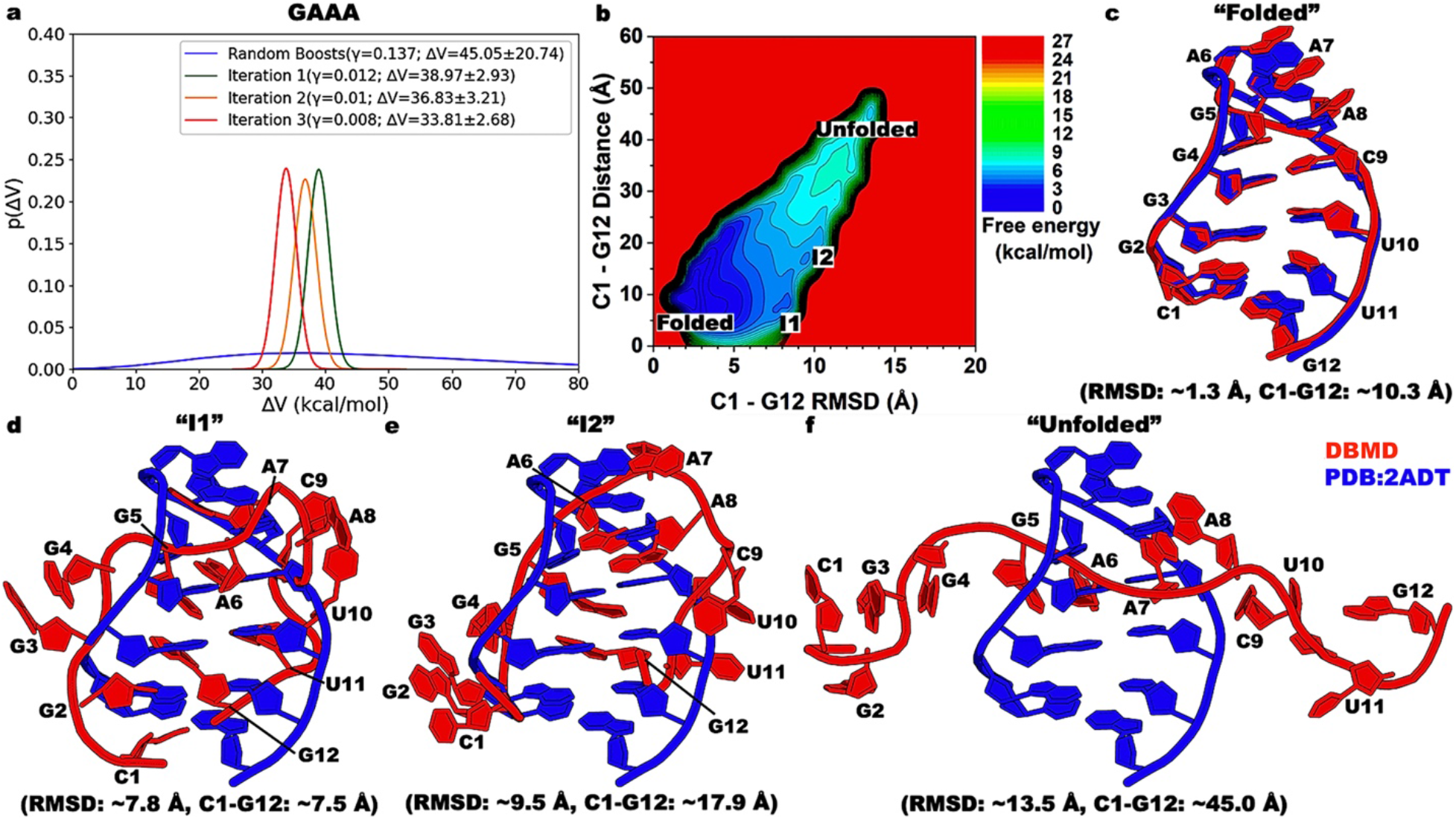
Folding of the 12-mer hairpin RNA with GAAA tetraloop in implicit solvent captured by DBMD. **(a)** Representative distributions of randomly generated dual boost potentials and DL-generated boost potentials iterated until *γ* < 0.01 from the potential energies collected from the pre-equilibration of the 12-mer hairpin RNA with GAAA tetraloop. The legends include the anharmonicity and average ± standard deviation of the dual boost potentials. **(b)** 2D PMF free energy profile of the heavy-atom RMSD of the 12-mer hairpin RNA relative to the 2ADT PDB and the COM distance between terminal nucleotides C1 and G12. The low-energy conformational states are labeled “Folded”, “I1”, “I2”, and “Unfolded”. **(c)** The “Folded” low-energy conformational state compared to the 2ADT PDB structure, for which the RMSD is ∼1.3 Å and the C1-G12 distance is ∼10.3 Å. **(d)** The intermediate “I1” low-energy conformational state compared to the 2ADT PDB structure, for which the RMSD is ∼7.8 Å and the C1-G12 distance is ∼7.5 Å. **(e)** The intermediate “I2” low-energy conformational state compared to the 2ADT PDB structure, for which the RMSD is ∼9.5 Å and the C1-G12 distance is ∼17.9 Å. **(f)** The “Unfolded” low-energy conformational state compared to the 2ADT PDB structure, for which the RMSD is ∼13.5 Å and the C1-G12 distance is ∼45.0 Å. The low-energy conformational states are colored red, and the 2ADT PDB structure is colored blue.

**Figure 6.**
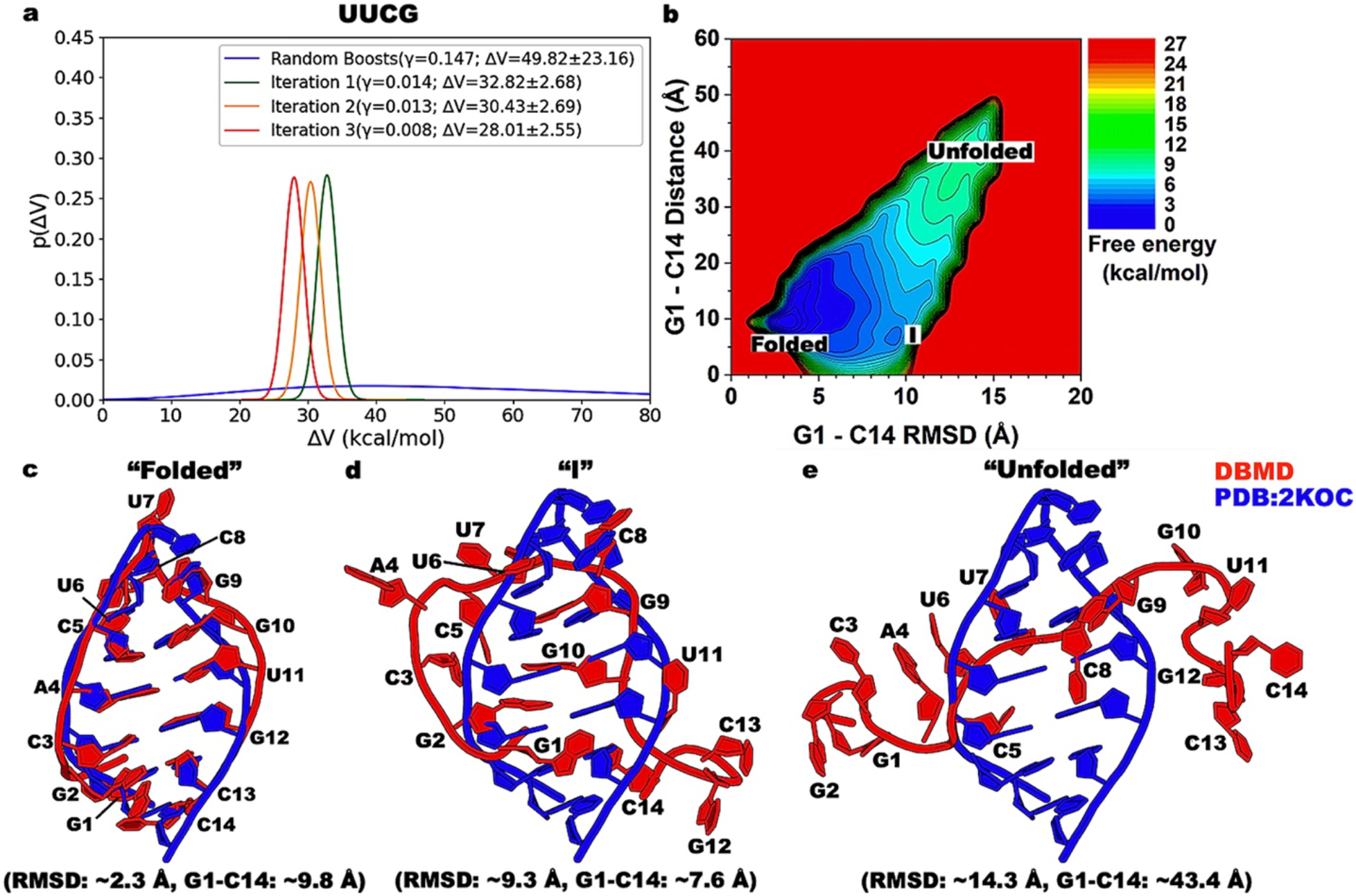
Folding of the 14-mer hairpin RNA with UUCG tetraloop in implicit solvent captured by DBMD. **(a)** Representative distributions of randomly generated dual boost potentials and DL-generated boost potentials iterated until *γ* < 0.01 from the potential energies collected from the pre-equilibration of the 14-mer hairpin RNA with UUCG tetraloop. The legends include the anharmonicity and average ± standard deviation of the dual boost potentials. **(b)** 2D PMF free energy profile of the heavy-atom RMSD of the 14-mer hairpin RNA relative to the 2KOC PDB and the COM distance between terminal nucleotides G1 and C14. The low-energy conformational states are labeled “Folded”, “I”, and “Unfolded”. **(c)** The “Folded” low-energy conformational state compared to the 2KOC PDB structure, for which the RMSD is ∼2.3 Å and the G1-C14 distance is ∼9.8 Å. **(d)** The intermediate “I” low-energy conformational state compared to the 2KOC PDB structure, for which the RMSD is ∼9.3 Å and the G1-C14 distance is ∼7.6 Å. **(e)** The “Unfolded” low-energy conformational state compared to the 2KOC PDB structure, for which the RMSD is ∼14.3 Å and the G1-C14 distance is ∼43.4 Å. The low-energy conformational states are colored red, and the 2KOC PDB structure is colored blue.

The time courses of the effective harmonic force constants (*k_0P_* and *k_0D_*) as well as the total and dihedral boost potential parameters (*V_min_*, *V_max_*, and *E*) during the equilibration of the hairpin RNAs with GCAA, GAAA, and UUCG tetraloop in implicit solvent are shown in **Figures S6-S8**. During the three rounds of 5ns DBMD equilibration of the GCAA RNA tetraloop system, the total effective harmonic force constant *k_0P_* decreased from 0.20 in round one to 0.10 in round two but increased to 0.17 in round three, while the dihedral effective harmonic force constant *k_0D_* decreased from 0.84 in round one to 0.56 in round two and 0.51 in round three (**Figure S6a**). The minimum total potential energy *V_minP_* fluctuated from −2867.79 kcal/mol in round one to −2930.46 kcal/mol in round two to −2871.63 kcal/mol in round three (**Figure S6b**). The maximum total potential energy *V_maxP_* also fluctuated between −2509.02 kcal/mol, −2366.52 kcal/mol, and −2458.10 kcal/mol among the three (**Figure S6b**). The minimum dihedral potential energy *V_minD_* fluctuated from 291.14 kcal/mol in round one to 326.13 kcal/mol in round two to 314.70 kcal/mol in round three, whereas the maximum dihedral potential energy *V_maxD_* decreased from 400.88 kcal/mol to 391.04 and 368.91 kcal/mol from round one to round three (**Figure S6c**). The reference total and dihedral potential energies *E_P_* and *E_D_* were mostly identical to the maximum total and dihedral potential energies *V_maxP_* and *V_maxD_*, except during round one for the *E_D_* (**Figure S6c**).

For the GAAA RNA tetraloop system, the total and dihedral effective harmonic force constants *k_0P_* and *k_0D_* decreased from 1.0 and 0.33 in round one to 0.15 and 0.98 in round two to 0.098 and 0.51 in round three (**Figure S7a**). The minimum total potential energy *V_minP_* decreased from −2621.20 kcal/mol in round one to −2746.68 kcal/mol in round two to −2799.21 kcal/mol in round three, whereas the maximum total potential energy *V_maxP_* fluctuated between −2380.18 kcal/mol, −2177.94 kcal/mol, and −2265.46 kcal/mol during the three rounds of DBMD equilibration (**Figure S7b**). The minimum dihedral potential energy *V_minD_* increased from 289.55 kcal/mol to 323.12 kcal/mol and 331.20 kcal/mol from round one to round three, while the maximum dihedral potential energy *V_maxD_* decreased from 396.04 kcal/mol to 387.99 and 380.68 kcal/mol from round one to three (**Figure S7c**). The reference total and dihedral potential energies were identical to the maximum total and dihedral energies.

For the UUCG RNA tetraloop system, the total and dihedral effective harmonic force constants *k_0P_* and *k_0D_* fluctuated between 0.12 and 1.0 in round one to 0.27 and 0.32 in round two to 0.19 and 0.46 in round three (**Figure S8a**). The minimum total potential energy *V_minP_* also fluctuated between −3360.51 kcal/mol, −3189.11 kcal/mol, and −3258.47 kcal/mol from round one to three of DBMD equilibration, whereas the maximum total potential energy *V_maxP_* increased from −2900.68 kcal/mol in round one to −2831.27 kcal/mol and −2733.89 kcal/mol in round two and three, respectively (**Figure S8b**). The minimum dihedral potential energy *V_minD_* fluctuated between 341.78 kcal/mol, 337.89 kcal/mol, and 327.41 kcal/mol from round one to three of DBMD equilibration, whereas the maximum dihedral potential energy *V_maxD_* decreased from 416.37 kcal/mol to 411.57 kcal/mol to 407.42 kcal/mol from round one to three (**Figure S8c**). The reference total and dihedral potential energies were the same as the maximum potential energies during the DBMD equilibration of the UUCG RNA tetraloop system.

Multiple independent 2µs DBMD simulations were performed on the hairpin RNAs with GCAA, GAAA, and UUCG tetraloops in implicit solvent, starting from their extended conformations (Figures 4-6). Remarkably, DBMD was able to capture multiple folding and unfolding events for all three hairpin RNAs within 2µs of simulations. In particular, a total of 18, 16, and 11 different stable folding-unfolding events were observed within 2µs DBMD simulations of the RNAs with GCAA, GAAA, and UUCG tetraloops, respectively (**Figures S9a-S11a**). The DBMD boost potentials exhibited Gaussian distributions, with low anharmonicity of 8.3 × 10^−3^, 3.9 × 10^−4^, and 2.9 × 10^−3^ in the GCAA, GAAA, and UUCG RNA tetraloop simulations (**Figures S9c, S10c,** and **S11c**). Furthermore, the boost potentials were recorded to be 37.0 ± 4.5 kcal/mol for the GCAA, 32.9 ± 3.1 kcal/mol for GAAA, and 27.6 ± 3.4 for UUCG system, given the different *η*_0_ and *η*_1_ used for the RNA systems.

The 2D PMF free energy profiles of the hairpin RNAs with tetraloops were calculated using the heavy-atom RMSDs of the whole RNAs relative to respective PDB structures (1ZIH^47^ for GCAA, 2ADT^48^ for GAAA, and 2KOC^49^ for UUCG) and the G1-U12, C1-G12, and G1-C14 center-of-mass (COM) distances as RCs. DBMD sampled three different low-energy conformational states, including “Folded”, intermediate “I”, and “Unfolded”, for the RNA with GCAA tetraloop (Figure 4b), four different low-energy conformational states, namely “Folded”, intermediate “I1” and “I2”, and “Unfolded”, for the GAAA tetraloop (Figure 5b), and three different low-energy conformational states, including “Folded”, intermediate “I”, and “Unfolded”, for the UUCG tetraloop (Figure 6b).

In the “Folded” low-energy conformational state of the 12-mer hairpin RNA with the GCAA tetraloop, the heavy-atom RMSD relative to the 1ZIH^47^ PDB structure was ∼1.1 Å, and the COM distance between terminal nucleotides G1 and U12 was ∼10.7 Å. This “Folded” low-energy conformational state was maintained by the Watson-Crick base pairs between nucleotides G2-C11, G3-C10, and C4-G9 and base stacking between nucleotides C6-A7-A8 of the GCAA tetraloop (Figure 4c). With transition from the “Folded” to the intermediate “I” state, most of the Watson-Crick base pairs distorted, with the side chains of nucleotides G9, C11, and U12 flipping out and exposing themselves to the solvent, while the base stacking between nucleotides C6-A7-A8 of the GCAA tetraloop was intact as observed in a conformation at ∼8.3Å heavy-atom RMSD relative to the 1ZIH^47^ PDB structure and ∼6.5Å G1-U12 COM distance (**Figure S12**). In the intermediate “I” low-energy conformational state, the RNA began extending, with nucleotides G1-C4 and C10-U12 extending in opposite directions. The base stacking between nucleotides C6-A7-A8 was mostly broken, with nucleotide A8 flipping out to base stack with nucleotide A9. In this state, the heavy-atom RMSD relative to the 1ZIH^47^ PDB structure was ∼9.2 Å, and the G1-U12 COM distance was ∼12.9 Å (Figure 4d). In the “Unfolded” low-energy conformational state, the RNA was completely stretched out, with a heavy-atom RMSD of ∼14.5 Å and G1-U12 COM distance of ∼48.3 Å (Figure 4e).

In the “Folded” low-energy conformational state of the 12-mer hairpin RNA with the GAAA tetraloop, the heavy-atom RMSD relative to the 2ADT^48^ PDB structure was ∼1.3 Å, and the COM distance between terminal nucleotides C1 and G12 was ∼10.3 Å. Similar to the GCAA system, this “Folded” state of the GAAA system was maintained by the Watson-Crick base pairs between nucleotides C1-G12 and G4-C9 as well as the base stacking between nucleotides A6-A7-A8 of the GAAA tetraloop (Figure 5c). The heavy-atom RMSD increased to ∼7.8 Å, whereas the C1-G12 COM distance decreased to ∼7.5 Å in the intermediate “I1” low-energy conformation. In this state, both the Watson-Crick base pairs and base stacking in the GAAA tetraloop were broken, with nucleotides G3, G4, A7, A8, C9, U10 flipping out and exposing to the solvent. However, base stacking was observed between nucleotides G5 and A6 of the GAAA tetraloop (Figure 5d). In the “I2” intermediate state, the heavy-atom RMSD the 2ADT^48^ PDB structure was ∼9.5 Å, and the COM distance between terminal nucleotides C1 and G12 was ∼17.9 Å. The 12-mer RNA was mostly distorted, with random base stacking formed between nucleotides G5-G12 and A6-A8. The side chains of the other nucleotides flipped out and exposed to the solvent (Figure 5e). In the “Unfolded” low-energy conformational state, the RNA was completely stretched out, with a heavy-atom RMSD of ∼13.5 Å and C1-G12 COM distance of ∼45.0 Å (Figure 5f).

The “Folded” low-energy conformational state of the 14-mer hairpin RNA with the UUCG tetraloop has a heavy-atom RMSD relative to the 2KOC^49^ PDB structure of ∼2.3 Å and COM distance between terminal nucleotides G1 and C14 of ∼9.8 Å. In this state, Watson-Crick base pairs were formed between nucleotides G2-C13, C3-G12, A4-U11, and C5-G10. However, unlike the GCAA and GCAA systems, no base stacking was observed between the nucleotides in the UUCG tetraloop (Figure 6c). In the “I” intermediate state, the heavy-atom RMSD increased to ∼9.3 Å and the G1-C14 COM distance decreased to ∼7.6 Å. The RNA was mostly distorted, with random base stacking formed between nucleotides G2 and G10. Most of the other nucleotides flipped out and exposed to the solvent (Figure 6d). In the “Unfolded” low-energy conformational state, the heavy-atom RMSD relative to the 2KOC^49^ PDB structure further increased to ∼14.3 Å and the COM distance between nucleotides G1 and C14 increased to ∼43.4 Å. The RNA was mostly stretched out (Figure 6e). Therefore, DBMD was able to capture repetitive folding and unfolding of RNA tetraloop structure in 2µs simulations, thereby enabling characterization of the RNA folding free energy landscapes.

## Discussion

In this work, we have developed DBMD, which generates boost potentials with Gaussian distribution using DL to reduce energy barriers and enhanced conformational sampling of biomolecules. Probabilistic Bayesian DL models are trained using potential energies of finished simulation frames to build the boost potentials that exhibit Gaussian distribution with anharmonicity *γ* < 0.01. We have demonstrated DBMD on the simulations of alanine dipeptide in explicit and implicit solvent and folding of the chignolin protein and hairpin RNAs with the GCAA, GAAA, and UUCG tetraloops. Overall, DBMD was able to greatly enhance conformational transitions and characterize the protein and RNA folding free energy landscapes.

DBMD captured multiple folding and unfolding events of chignolin within 300 ns of simulations (**Figure S5a**). Compared to previous cMD performed with Anton^56^ and aMD^57^ simulations of chignolin folding, DBMD sped up the folding-unfolding transition by ∼69,489 and 1.35 times, respectively. Furthermore, compared to previous 300ns simulations performed with GaMD in AMBER^27^ and NAMD^29^, DBMD accelerated the folding-unfolding transition by 6 times, while still providing a 2D free energy profile of the C_α_-atom RMSD and Rg of residues Y2-W9 with high degrees of similarity^27, 29^. In particular, DBMD sampled all three low-energy conformational states (“Folded”, intermediate “I”, and “Unfolded”) as GaMD in AMBER^27^ and the two low-energy conformations (“Folded” and “I”) as GaMD in NAMD^29^ (Figure 3). Moreover, the folding mechanism uncovered by DBMD was relatively similar to that by GaMD in AMBER^27^. Starting from the extended conformation of the “Unfolded” state (Figure 3e), the terminal residues of chignolin was brought closer due to the interactions between residues P4 and G7 in the intermediate “I” state (Figure 3d). With transition from the intermediate “I” to the “Folded” state, antiparallel β-sheets were formed between residues G1-D3 and G7-G10, with the hydrophilic side chains of residues D3, E5, T6, and T8 exposed to the solvent (Figure 3c).

For the simulations of the hairpin RNAs with GCAA, GAAA, and UUCG tetraloops, the total number of folding and unfolding events captured by AIMBD simulations reduced from the GCAA to GAAA to UUCG simulation system, which was in good agreement with previous studies by Tan et al.^46^ and Chen et al.^58^. This also demonstrated the importance of the base stacking within the tetraloop for RNA folding. In particular, while nucleotides C6-A7-A8 of the GCAA tetraloop and A6-A7-A8 of the GAAA tetraloop base-stacked in their respective “Folded” low-energy conformations, no base stacking was observed within the “Folded” hairpin RNA with UUCG tetraloop (Figures 4c-6c). Furthermore, the folding mechanisms uncovered by DBMD were similar among the hairpin RNAs with GCAA, GAAA, and UUCG tetraloop (Figures 4-6). Starting from the extended conformation in the “Unfolded” low-energy conformational states (Figures 4e, 5f, and 6e), Watson-Crick base pairs began to form from terminal nucleotides towards the cores and tetraloops of the RNAs. Finally, base stacking between the nucleotides of the tetraloops were formed to enable the stable folding of the hairpin RNAs (Figures 4c and 5c). This general mechanism of RNA folding showed high degrees of similarity to the previous study by Chen et al.^58^, even though they used shorter RNA strands, a different force field parameter set, and a different solvation model.

In conclusion, we have developed DBMD, a DL-based enhanced sampling technique that allows for accurate energetic reweighting and enhanced sampling of biomolecular systems. DBMD is available with open source in OpenMM at https://github.com/MiaoLab20/DBMD/. As demonstrated on the model systems, DBMD captured multiple dihedral transitions of alanine dipeptide as well as folding-unfolding events of the chignolin protein and hairpin RNAs with tetraloops within relatively short simulation lengths. DBMD is expected to facilitate the simulations and free energy calculations of a wide range of biomolecules.

## Author Contributions

H.N.D. performed research, analyzed data, and wrote the manuscript. Y.M. supervised the project, interpreted data, and wrote the manuscript. All authors contributed towards the final version of the manuscript.

## Acknowledgements

We thank Matthew Copeland and Dr. Jinan Wang for the valuable discussions. This work used supercomputing resources with allocation award TG-MCB180049 through the Advanced Cyberinfrastructure Coordination Ecosystem: Services & Support (ACCESS) program, which is supported by National Science Foundation grants #2138259, #2138286, #2138307, #2137603, and #2138296, and project M2874 through the National Energy Research Scientific Computing Center (NERSC), which is a U.S. Department of Energy Office of Science User Facility operated under Contract No. DE-AC02-05CH11231, and the Research Computing Cluster and BigJay Cluster funded through NSF Grant MRI-2117449 at the University of Kansas. This work was supported by the National Institutes of Health (R01GM132572).

