## Supporting Information for "Deep Boosted Molecular Dynamics (DBMD): Accelerating molecular simulations with Gaussian boost potentials generated using probabilistic Bayesian deep neural network"

**Hung N. Do[1] and Yinglong Miao[1,*]**

[1]Center for Computational Biology and Department of Molecular Biosciences, University of Kansas, Lawrence, Kansas 66047

**Algorithm of Deep Boosted Molecular Dynamics (DBMD)**

```
DBMD {
    // Stage 1: Conventional molecular dynamics
    For i = 1, ..., conventional_md_steps:
        If (i == conventional_md_steps):
            Vmin = min(V1, V2, ..., Vi)
            Vmax = max(V1, V2, ..., Vi)
    End

    // Stage 2: Pre-equilibration
    If (simulation_type == "explicit"):
        k0P = 1.0
        k0D = 1.0
    Else: // (simulation_type == "protein.implicit" or "RNA.implicit")
        k0P = 0.05
        k0D = 1.0
    Set refE_factor
    For i = 1, ..., pre_equilibration_steps:
        Record V
        E = Vmin + (Vmax – Vmin) / k0
        If (E > Vmax + refE_factor*abs(Vmax)):
            E = Vmax
        If (V < E):
            ΔV = (1/2)*k0*(E – V)^2 / (Vmax – Vmin)
            V = V + ΔV
        Vmin = min(V, Vmin)
        Vmax = max(V, Vmax)
    End

    // Stage 3: Equilibration
    // Deep Learning model
    Set mu, sigma       // default mu = 0.0, sigma = 1.0 for standard normal distribution
    Define PriorModel:
        PriorModel = Sequential {
            DistributionLambda(MultivariateNormalDiag(loc=mu*ones, scale_diag=sigma*ones)
            }
    Define PosteriorModel:
        PosteriorModel = Sequential {
            VariableLayer(MultivariateNormalTriL)
            MultivariateNormalTriL
```

```
        }
    Define BayesianNeuralNetworkModel:
        If (simulation_type == “explicit”):
            L2 = 1
        Else: // (simulation_type == “protein.implicit” or “RNA.implicit”)
            L2 = 3
        BayesianNeuralNetworkModel = Sequential {
            DenseVariational(64, input_dim = 1,
                            prior_function = PriorModel, posterior_function = PosteriorModel,
                            activation = “sigmoid”)
            For _ = 1, ..., L2:
                DenseVariational(IndependentNormal(1),
                            prior_function = PriorModel, posterior_function = PosteriorModel)
            End
            IndependentNormal(1)     // output layer
            }
        Compile(loss = Kullback-LeiberDivergence, optimizer = Adam(learning_rate=0.0003)

    // Deep Learning
    Collect {V1, V2, ..., VM}, Vmin, Vmax from the M-step pre-equilibration
    For i = 1, ..., M:
        k0 = random(0, 1]
        E = Vmin + (Vmax – Vmin) / k0
        ΔV = (1/2)*k0*(E – V)^2 / (Vmax – Vmin)
    End
    Collect {ΔV1, ΔV2, ..., ΔVM}
    While (anharmocity(ΔV) >= 0.01):
        training_set, validation_set = train_test_split({V1, V2, ..., VM}, {ΔV1, ΔV2, ..., ΔVM}, test_size = 0.2)
        BayesianNeuralNetworkModel.fit(training_set, epochs = 100, batch_size = 100, validation_set)
    ΔVM = BayesianNeuralNetworkModel.predict(VM)
    k0 = ((sqrt(2*ΔVM*(Vmax – Vmin)) – sqrt(2*ΔVM*(Vmax – Vmin) – 4*(Vmin – VM)*(Vmax – Vmin))) / (2*(Vmin – VM)))^2
    E = Vmin + (Vmax – Vmin) / k0
    If ((k0 > 1.0) or (E > Vmax + refE_factor*abs(Vmax))):
        E = Vmax
        k0 = (2*ΔVM*(Vmax – Vmin)) / (E – VM)^2
        k0 = min(1.0, k0)
    Collect Vmin, Vmax, k0 for equilibration
```

```
// Muti-round equilibration
For i = 1, ..., equilibration_rounds:
    For j = 1, ..., equilibration_steps_per_round:
        Record V
        E = Vmin + (Vmax – Vmin) / k0
        If (E > Vmax + refE_factor*abs(Vmax)):
            E = Vmax
        If (V < E):
            ΔV = (1/2)*k0*(E – V)^2 / (Vmax – Vmin)
            V = V + ΔV
        Vmin = min(V, Vmin)
        Vmax = max(V, Vmax)
    Collect {V1, V2, ..., VN}, Vmin, Vmax from the N-step equilibration round
    For i = 1, ..., N:
        k0 = random(0, 1]
        E = Vmin + (Vmax – Vmin) / k0
        ΔV = (1/2)*k0*(E – V)^2 / (Vmax – Vmin)
    End
    Collect {ΔV1, ΔV2, ..., ΔVN}
    While (anharmocity(ΔV) >= 0.01):
        training_set, validation_set = train_test_split({V1, V2, ..., VN}, {ΔV1, ΔV2, ..., ΔVN}, test_size = 0.2)
        BayesianNeuralNetworkModel.fit(training_set, epochs = 100, batch_size = 100, validation_set)
    ΔVN = BayesianNeuralNetworkModel.predict(VN)
    k0 = ((sqrt(2*ΔVN*(Vmax – Vmin)) – sqrt(2*ΔVN*(Vmax – Vmin) – 4*(Vmin – VN)*(Vmax – Vmin))) / (2*(Vmin – VN)))^2
    E = Vmin + (Vmax – Vmin) / k0
    If ((k0 > 1.0) or (E > Vmax + refE_factor*abs(Vmax))):
        E = Vmax
        k0 = (2*ΔVN*(Vmax – Vmin)) / (E – VN)^2
        k0 = min(1.0, k0)
    Collect Vmin, Vmax, k0 for the next equilibration round
    If (i = equilibration_rounds):
        Collect Vmin, Vmax, k0 for production
    End
End

// Stage 4: Production
```

```
    For i = 1, ..., production_steps:
        E = Vmin + (Vmax – Vmin) / k0
            If (E > Vmax + refE_factor*abs(Vmax)):
                E = Vmax
            If (V < E):
                ΔV = (1/2)*k0*(E – V)^2 / (Vmax – Vmin)
                V = V + ΔV
    End
}
```

**Example Input Python File for DBMD Simulation with OpenMM**

```
parmFile = "dip.top"            # topology file
crdFile = "dip.crd"             # coordinate file
simType = "explicit"            # "explicit", "protein.implicit", "RNA.implicit"

temperature = 300       # simulation temperature

ntcmd = 1000000         # number_of_conventional_MD_steps
cmdRestartFreq = 100            # conventional_MD_restart_frequency

ncycebprepstart, ncycebprepend = 0, 1           # pre_equilibration_start and _end_round_indices
ntebpreppercyc = 1000000            # number_of_pre_equilibration_steps_per_round
ebprepRestartFreq = 100             # pre_equilibration_restart_frequency

ncycebstart, ncycebend = 0, 1               # equilibration_start and _end_round_indices
ntebpercyc = 1000000            # number_of_equilibration_steps_per_round
ebRestartFreq = 100             # equilibration_restart_frequency

ncycprodstart, ncycprodend = 0, 3           # production_start and _end_round_indices
ntprodpercyc = 5000000              # number_of_production_steps_per_round
prodRestartFreq = 10                # production_restart_frequency

refEP_factor, refED_factor = 0.0, 0.0       # reference energy factor; value between 0 and 1
```

**Description of DBMD Parameters in Input Python File for Simulation with OpenMM**

| | |
|---|---|
| parmFile | Path to the system topology file, usually a **parm7** or **prmtop** file. |
| crdFile | Path to the system coordinate file, usually a **crd** or **rst7** file. |
| simType | Solvation model and type of biomolecules simulated. Three variables are accepted:<br>• "explicit" (explicit-solvent simulations),<br>• "protein.implicit" (implicit-solvent simulations of proteins),<br>• "RNA.implicit" (implicit-solvent simulations of RNAs). |
| temperature | Simulation temperature in Kelvins (K). |
| ntcmd | Number of simulation steps in the conventional MD (cMD) stage. |
| cmdRestartFreq | Number of simulation steps per which the cMD trajectories and simulation checkpoints are outputted. |
| ncycebprepstart | DBMD pre-equilibration can be carried out in multiple rounds. The index of the starting round in DBMD pre-equilibration simulation. |
| ncycebprepend | The index of the final round in DBMD pre-equilibration simulation. |
| ntebpreppercyc | Number of simulation steps per round in the DBMD pre-equilibration. |
| ebprepRestartFreq | Number of simulation steps per which the pre-equilibration trajectories and simulation checkpoints are outputted. |
| ncycebstart | DBMD equilibration can be carried out in multiple rounds. The index of the starting round in DBMD equilibration simulation. |
| ncycebend | The index of the final round in DBMD equilibration simulation. |
| ntebpercyc | Number of simulation steps per round in the DBMD equilibration. |
| ebRestartFreq | Number of simulation steps per which the equilibration trajectories and simulation checkpoints are outputted. |
| ncycprodstart | DBMD production can be carried out in multiple rounds. The index of the starting round in DBMD production simulation. |
| ncycprodend | The index of the final round in DBMD production simulation. |
| ntprodpercyc | Number of simulation steps per round in the DBMD production. |
| prodRestartFreq | Number of simulation steps per which the production trajectories and simulation checkpoints are outputted. |
| refEP_factor | The reference total potential energy factor ($\eta_P$) for applying total boost potentials. This parameter, valued between 0 and 1, is introduced to avoid exceedingly large reference potential energy. The upper limit for the reference total potential energy $E_P$ is $V_{maxP} + \eta_P \lvert V_{maxP} \rvert$. |
| refED_factor | The reference dihedral potential energy factor ($\eta_D$) for applying dihedral boost potentials. This parameter, valued between 0 and 1, is introduced to avoid exceedingly large reference potential energy. The upper limit for the reference dihedral potential energy $E_D$ is $V_{maxD} + \eta_D \lvert V_{maxD} \rvert$. |

**Figure S1. (a-d)** Time courses of the effective harmonic force constants ($k_{0P}$ and $k_{0D}$) **(a)**, total **(b)** and dihedral **(c)** boost potential parameters ($V_{min}$, $V_{max}$, and $E$), and Phi dihedral **(d)** of alanine dipeptide calculated from one round of 2ns DBMD equilibration in explicit solvent. **(e-h)** Time courses of the effective harmonic force constants ($k_{0P}$ and $k_{0D}$) **(e)**, total **(f)** and dihedral **(g)** boost potential parameters ($V_{min}$, $V_{max}$, and $E$), and Phi (Φ) dihedral **(h)** of alanine dipeptide calculated from one round of 2ns DBMD equilibration in implicit solvent.

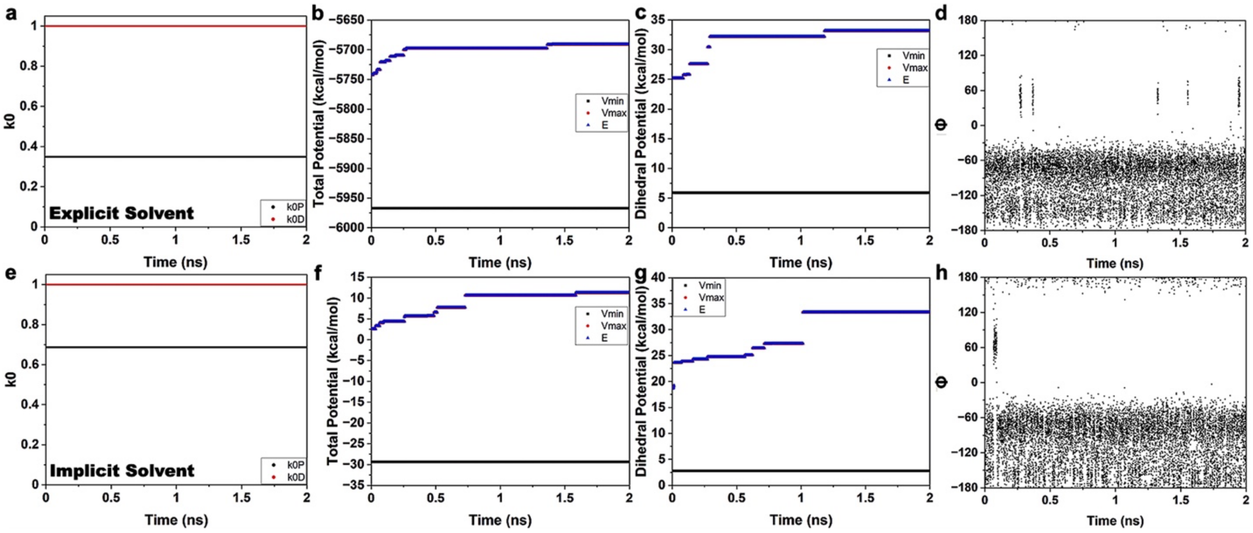

**Figure S2. (a-d)** Time courses of the Phi (Φ) dihedral of alanine dipeptide calculated from three 30ns DBMD simulations **(a-c)** and one 1μs cMD simulation **(d)** in explicit solvent. **(e-h)** Time courses of the Phi (Φ) dihedral of alanine dipeptide calculated from three 30ns DBMD simulations **(e-g)** and one 1μs cMD simulation **(h)** in implicit solvent.

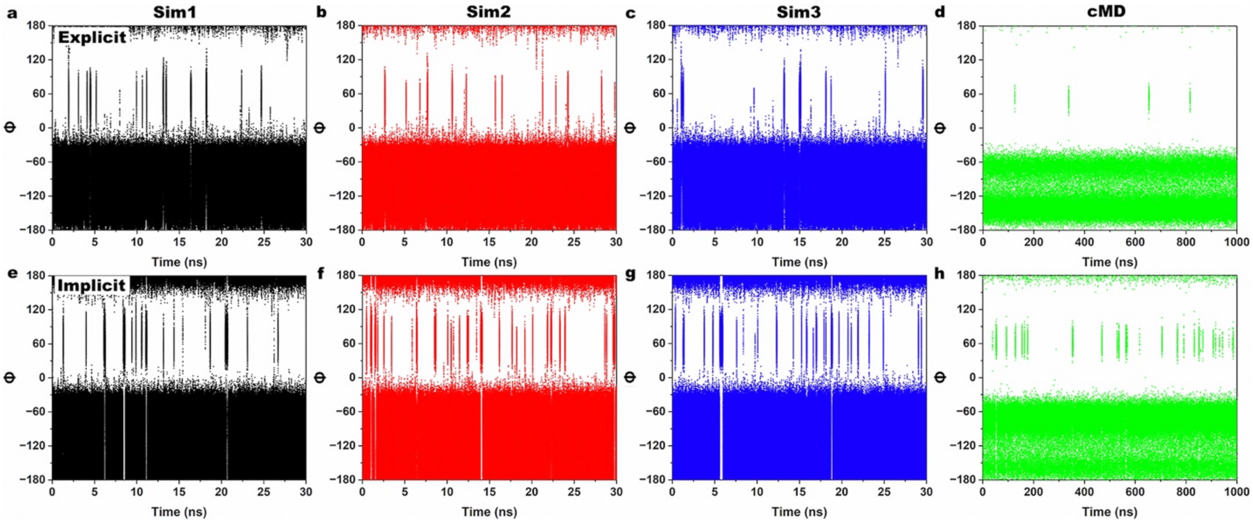

**Figure S3.** **(a-b)** Distributions of the boost potentials $\Delta V$ applied in the DBMD simulations of alanine dipeptide in explicit solvent **(a)** and implicit solvent **(b)**. **(c-f)** Potential of mean force (PMF) free energy profiles of the $\Phi$ **(c-d)** and $\Psi$ **(e-f)** dihedrals of alanine dipeptide calculated from three 30ns DBMD simulations compared to 1μs cMD simulations in explicit solvent **(c, e)** and implicit solvent **(d, f)**.

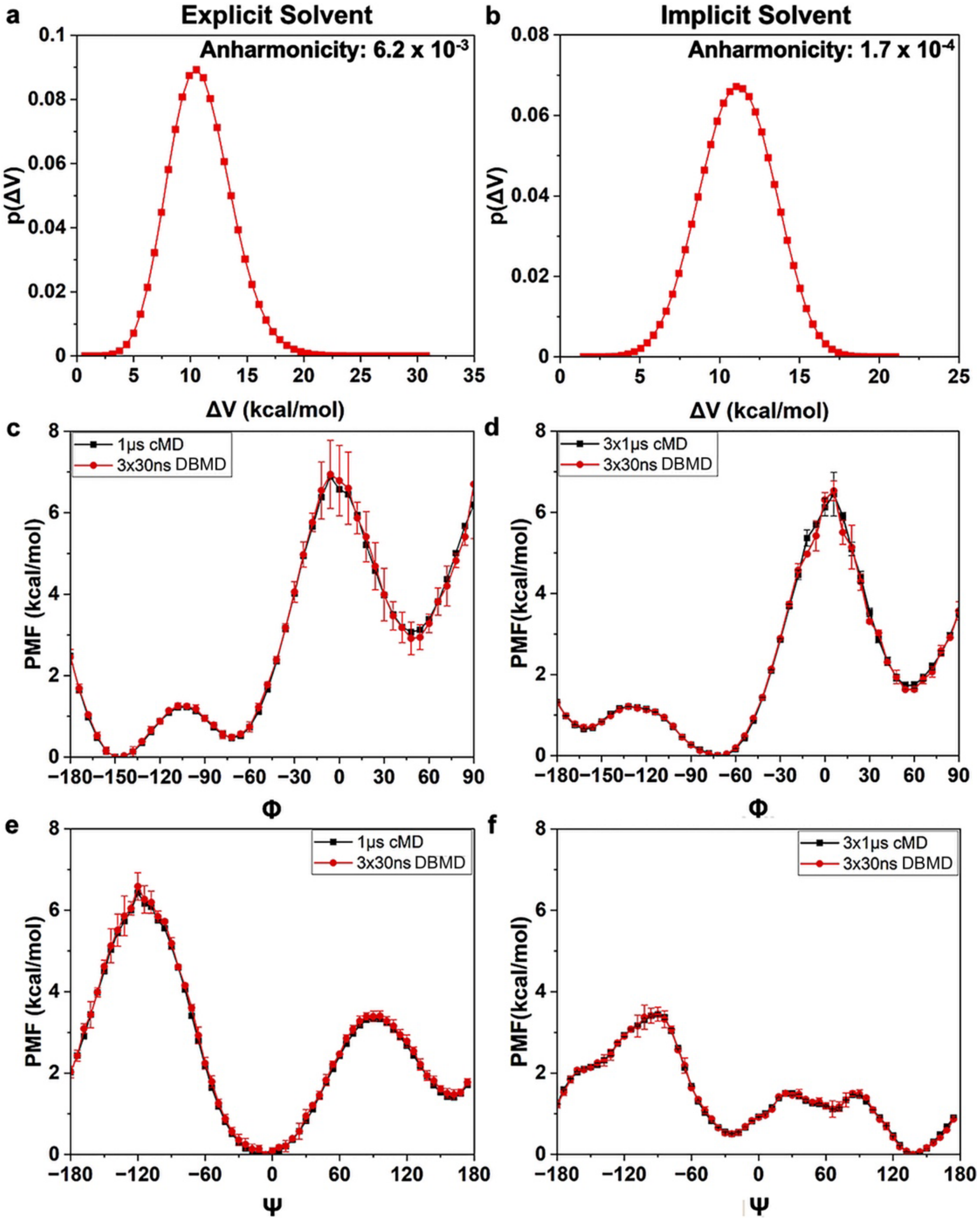

**Figure S4.** Time courses of the effective harmonic force constants ($k_{0P}$ and $k_{0D}$) **(a)**, total **(b)** and dihedral **(c)** boost potential parameters ($V_{min}$, $V_{max}$, and $E$), and $C_{\alpha}$-atom RMSD of residues Y2-W9 of chignolin relative to the 1UAO PDB **(d)** calculated from two rounds (*R1* and *R2*) of 5ns DBMD equilibration of chignolin in explicit solvent.

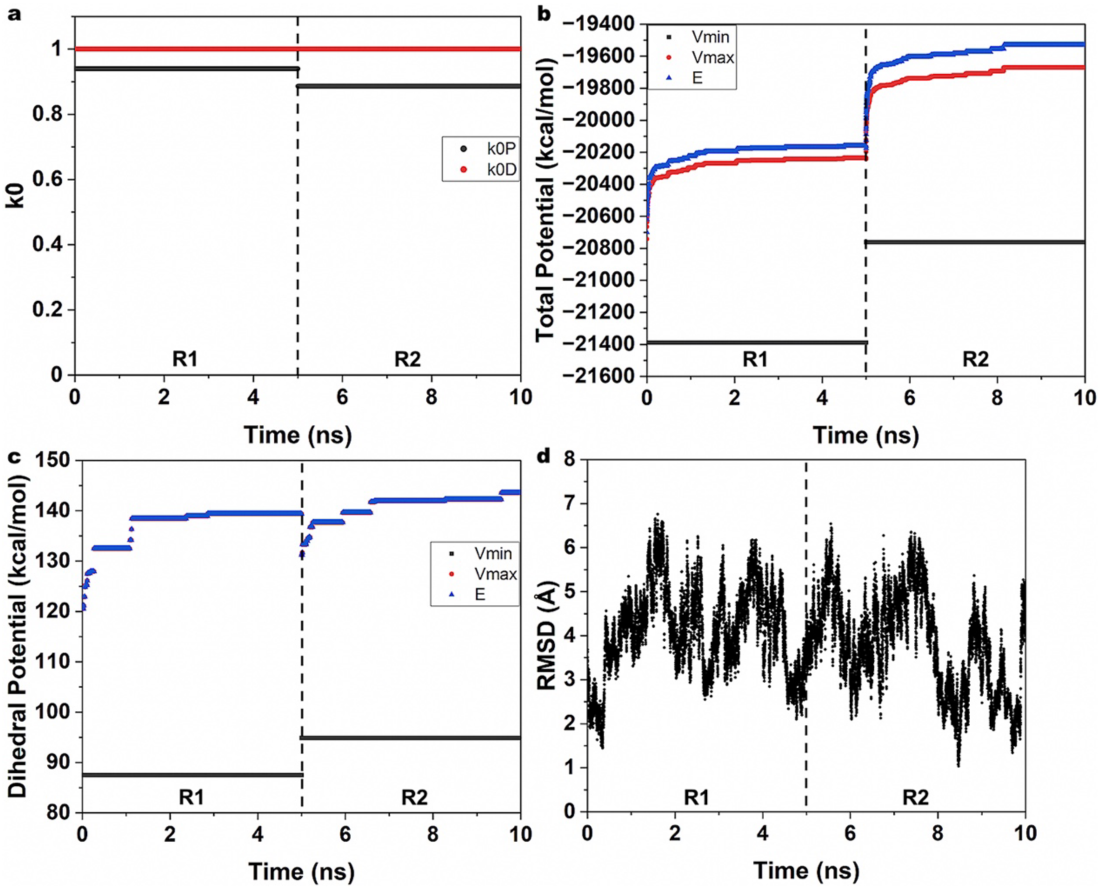

**Figure S5. (a-b)** Time courses of the $C_\alpha$-atom RMSD of residues Y2-W9 of chignolin relative to the 1UAO PDB **(a)** and $C_\alpha$-atom Rg of residues Y2-W9 **(b)** calculated from three 300ns DBMD simulations of chignolin in explicit solvent. **(c)** Distribution of the boost potentials $\Delta V$ applied in the DBMD simulations of chignolin in explicit solvent.

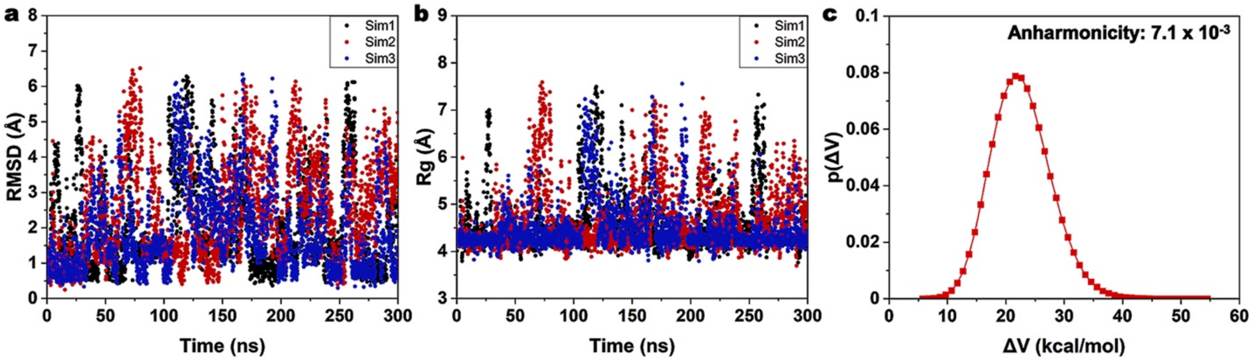

**Figure S6.** Time courses of the effective harmonic force constants ($k_{0P}$ and $k_{0D}$) **(a)**, total **(b)** and dihedral **(c)** boost potential parameters ($V_{min}$, $V_{max}$, and $E$), and heavy-atom RMSD of the hairpin RNA relative to the 1ZIH PDB **(d)** calculated from three rounds (*R1*, *R2*, and *R3*) of 5ns DBMD equilibration of the 12-mer hairpin RNA with GCAA tetraloop in implicit solvent.

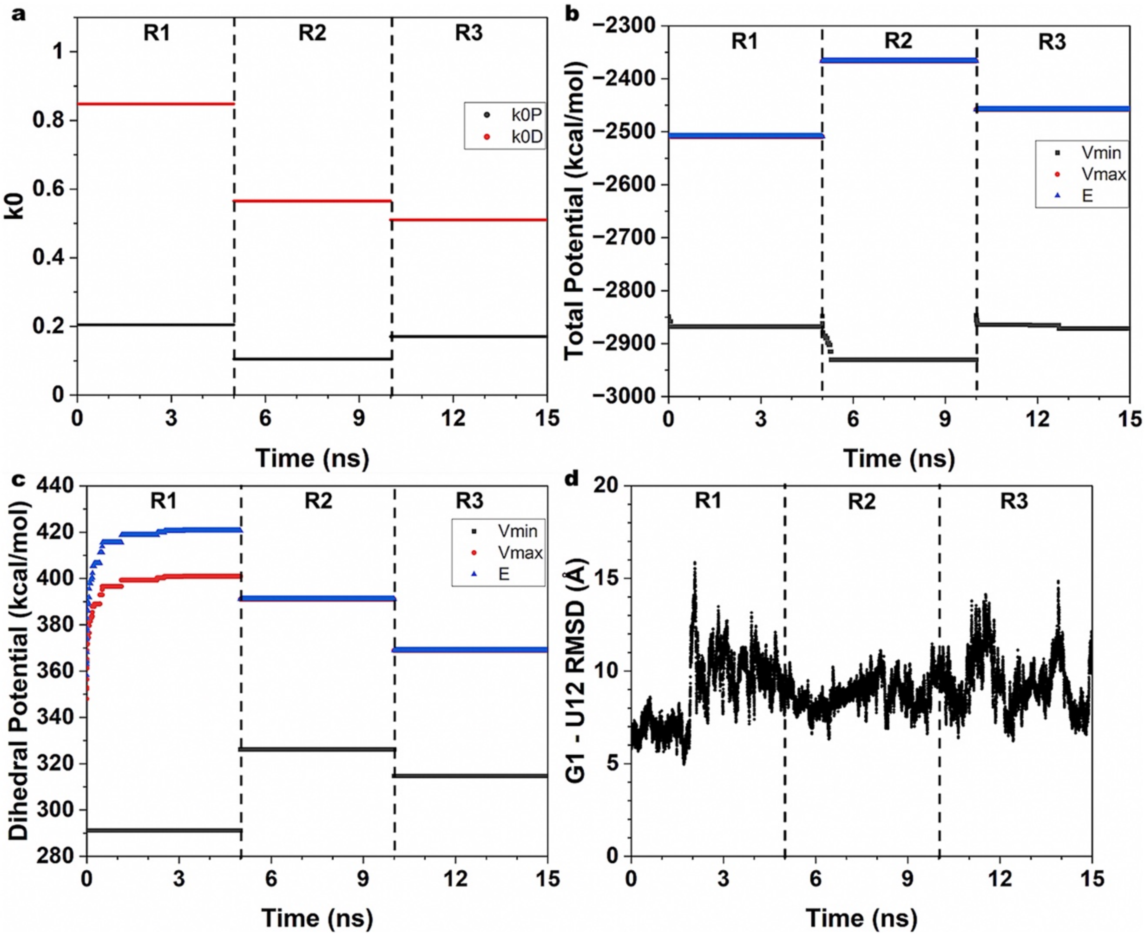

**Figure S7.** Time courses of the effective harmonic force constants ($k_{0P}$ and $k_{0D}$) **(a)**, total **(b)** and dihedral **(c)** boost potential parameters ($V_{min}$, $V_{max}$, and $E$), and heavy-atom RMSD of the hairpin RNA relative to the 2ADT PDB **(d)** calculated from three rounds (*R1*, *R2*, and *R3*) of 5ns DBMD equilibration of the 12-mer hairpin RNA with GAAA tetraloop in implicit solvent.

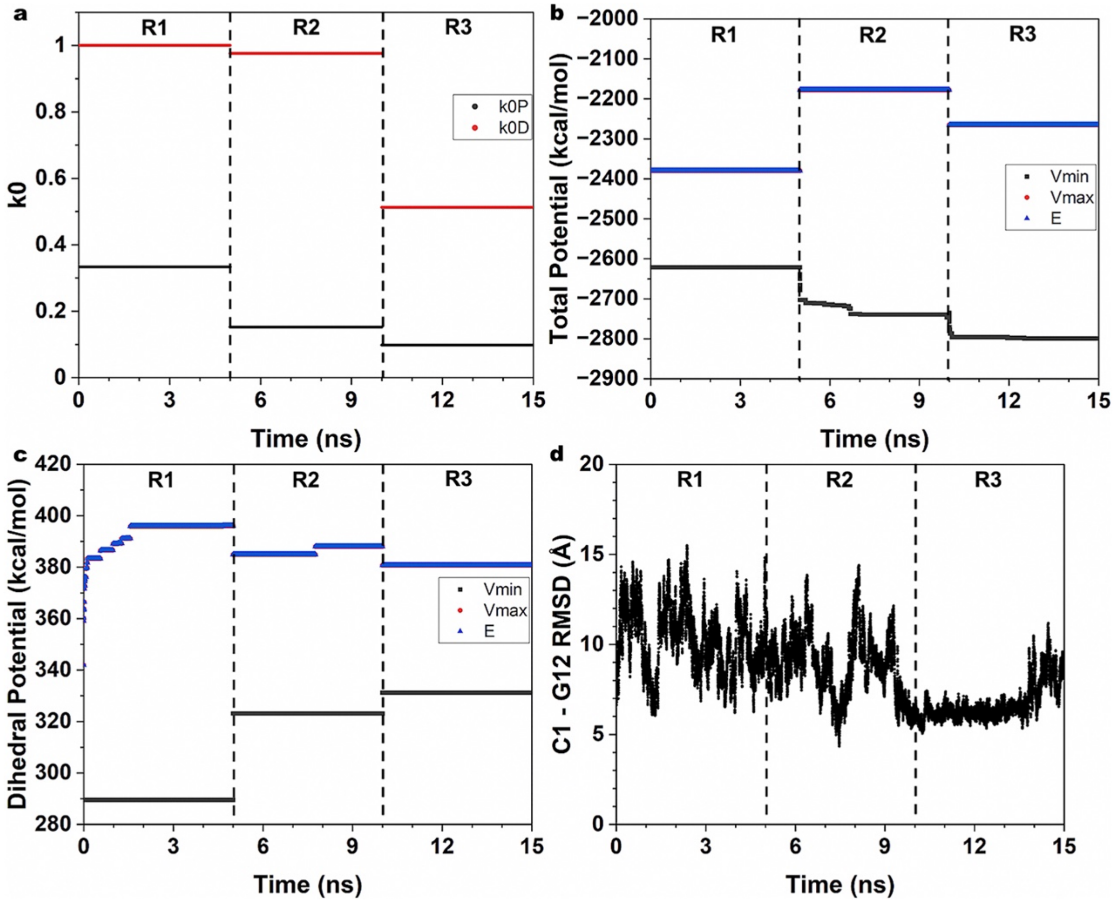

**Figure S8.** Time courses of the effective harmonic force constants ($k_{0P}$ and $k_{0D}$) **(a)**, total **(b)** and dihedral **(c)** boost potential parameters ($V_{min}$, $V_{max}$, and $E$), and heavy-atom RMSD of the hairpin RNA relative to the 2KOC PDB **(d)** calculated from three rounds (*R1*, *R2*, and *R3*) of 5ns DBMD equilibration of the 14-mer hairpin RNA with UUCG tetraloop in implicit solvent.

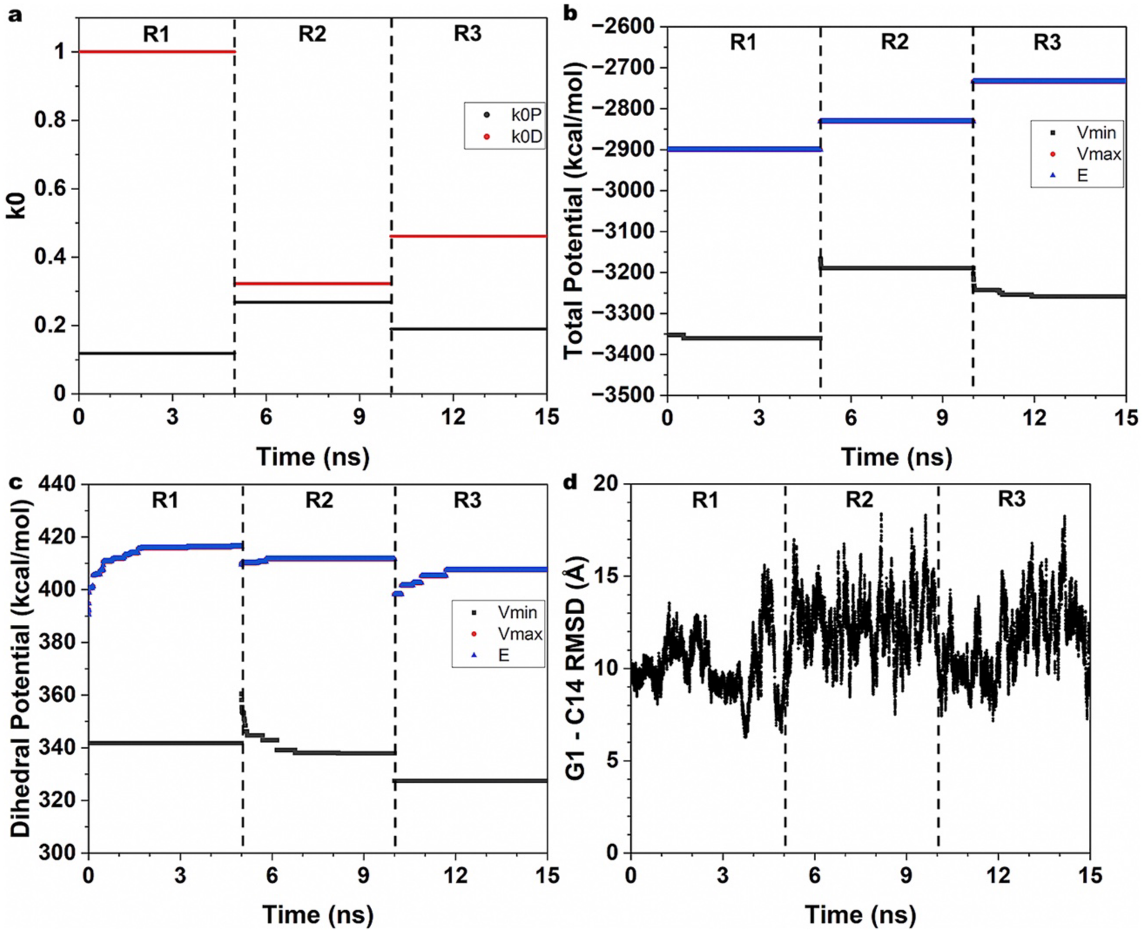

**Figure S9. (a-b)** Time courses of the heavy-atom RMSD of the 12-mer hairpin RNA with GCAA tetraloop relative to the 1ZIH PDB **(a)** and the COM distance between terminal nucleotides G1 and U12 **(b)** calculated from three 2000ns DBMD simulations of the 12-mer hairpin RNA with GCAA tetraloop in implicit solvent. **(c)** Distribution of the boost potentials $\Delta V$ applied in the DBMD simulations of the 12-mer hairpin RNA with GCAA tetraloop in implicit solvent.

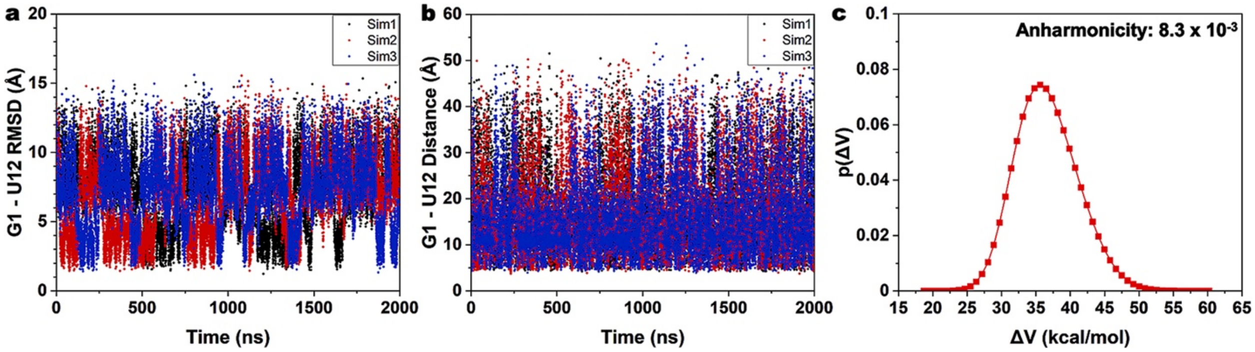

**Figure S10. (a-b)** Time courses of the heavy-atom RMSD of the 12-mer hairpin RNA with GAAA tetraloop relative to the 2ADT PDB **(a)** and the COM distance between terminal nucleotides C1 and G12 **(b)** calculated from three 2000ns DBMD simulations of the 12-mer hairpin RNA with GAAA tetraloop in implicit solvent. **(c)** Distribution of the boost potentials $\Delta V$ applied in the DBMD simulations of the 12-mer hairpin RNA with GAAA tetraloop in implicit solvent.

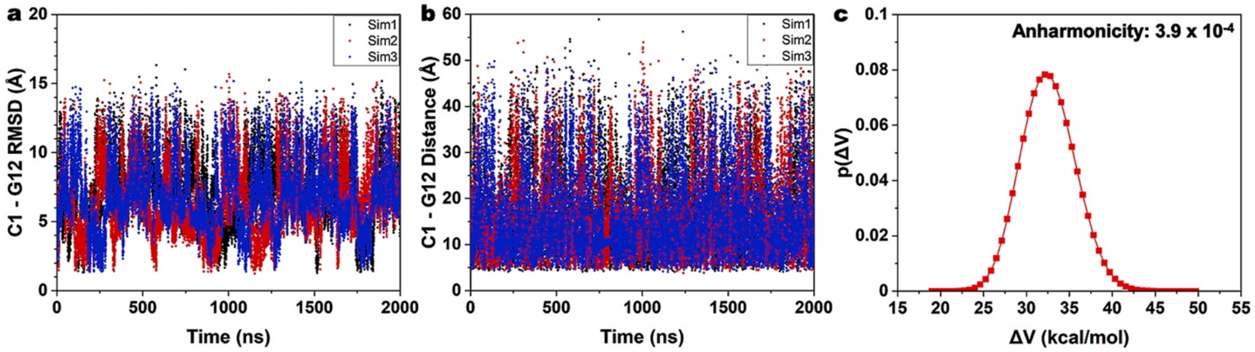

**Figure S11. (a-b)** Time courses of the heavy-atom RMSD of the 14-mer hairpin RNA with UUCG tetraloop relative to the 2KOC PDB **(a)** and the COM distance between terminal nucleotides G1 and C14 **(b)** calculated from four 2000ns DBMD simulations of the 14-mer hairpin RNA with UUCG tetraloop in implicit solvent. **(c)** Distribution of the boost potentials $\Delta V$ applied in the DBMD simulations of the 14-mer hairpin RNA with UUCG tetraloop in implicit solvent.

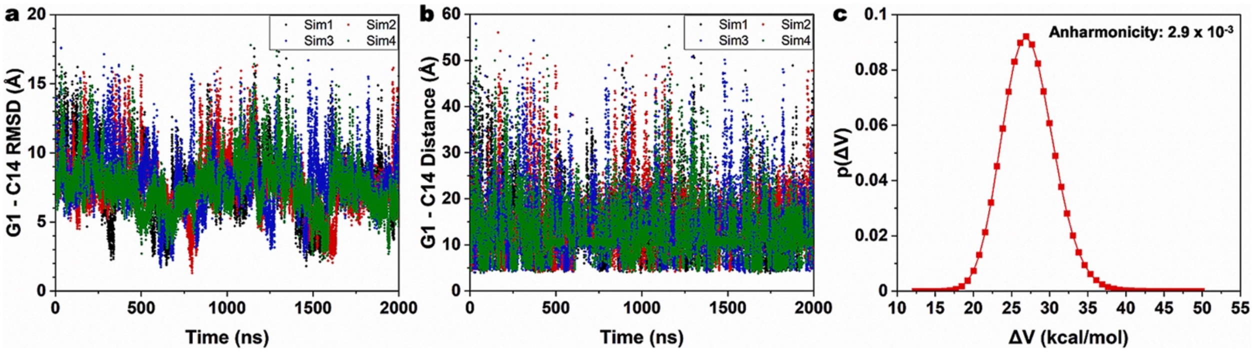

**Figure S12.** Conformation of the 12-mer hairpin RNA with the GCAA tetraloop at (~8.3 Å, ~6.5 Å) of the heavy-atom RMSD relative to the 1ZIH[47] PDB structure and G1-U12 COM distance.

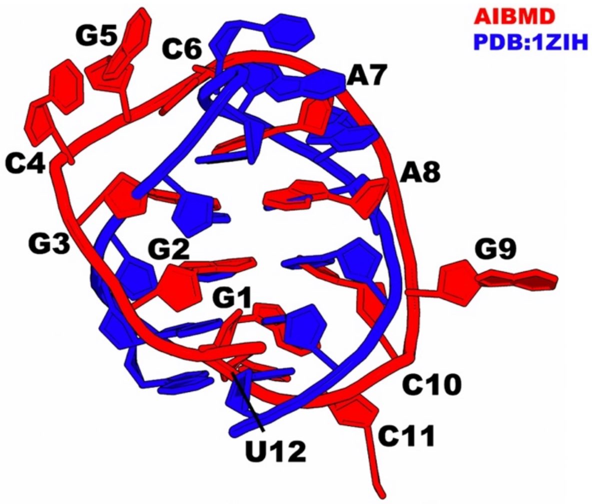


(RMSD: ~8.3 Å, G1-U12: ~6.5 Å)
